# Recombinant NAGLU-IGF2 prevents physical and neurological disease and improves survival in Sanfilippo B syndrome

**DOI:** 10.1101/2021.08.06.455469

**Authors:** Steven Q. Le, Shih-hsin Kan, Marie S. Roberts, Joshua T. Dearborn, Feng Wang, Shan Li, Elizabeth M. Snella, Jackie K. Jens, Bethann N. Valentine, Hemanth R. Nelvagal, Alexander Sorensen, Keerthana Chintalapati, Kevin Ohlemiller, Carole Vogler, Jonathan D. Cooper, Tsui-Fen Chou, N. Matthew Ellinwood, Jodi D. Smith, Mark S. Sands, Patricia I. Dickson

**Affiliations:** Washington University School of Medicine, St. Louis, Missouri, USA; CHOC Research Institute, Orange, California, USA; California Institute of Technology, Pasadena, California, USA; Iowa State University, Ames, Iowa, USA; Saint Louis University School of Medicine, St. Louis, Missouri, USA

**Keywords:** mucopolysaccharidosis, enzyme replacement therapy, glycosaminoglycan, proteomics, lysosomal storage disease

## Abstract

Recombinant human alpha-*N*-acetylglucosaminidase-insulin-like growth factor-2 (rhNAGLU-IGF2) is an investigational enzyme replacement therapy for Sanfilippo B, a lysosomal storage disease. Because recombinant human NAGLU (rhNAGLU) is poorly mannose 6-phosphorylated, we generated a fusion protein of NAGLU with IGF2 to permit its binding to the cation-independent mannose 6-phosphate receptor. We previously administered rhNAGLU-IGF2 intracerebroventricularly to Sanfilippo B mice, and demonstrated therapeutic restoration of NAGLU, normalization of lysosomal storage, and improvement in markers of neurodegeneration and inflammation. Here, we studied repeated intracerebroventricular rhNAGLU-IGF2 delivery in both murine and canine Sanfilippo B to determine potential effects on their behavioral phenotypes and survival. Treated mice showed improvement in disease markers such as heparan sulfate glycosaminoglycans, beta-hexosaminidase, microglial activation, and lysosomal-associated membrane protein-1. Sanfilippo B mice treated with rhNAGLU-IGF2 displayed partial normalization of their stretch attend postures, a defined fear pose in mice (p<0.001). We found an improved rotarod performance in Sanfilippo B mice treated with rhNAGLU-IGF2 compared to vehicle-treated Sanfilippo B mice (p=0.002). We also found a 61% increase in survival in Sanfilippo B mice treated with rhNAGLU-IGF2 (mean 53w, median 48w) compared to vehicle-treated Sanfilippo B mice (mean 33w, median 37w; p<0.001). In canine Sanfilippo B, we found that rhNAGLU-IGF2 administered into cerebrospinal fluid normalized HS and beta-hexosaminidase activity in gray and white matter brain regions. Proteomic analysis of cerebral cortex showed restoration of protein expression levels in pathways relevant to cognitive, synaptic, and lysosomal functions. These data suggest that treatment with rhNAGLU-IGF2 may improve the phenotype of Sanfilippo B disease.

## Introduction

Lysosomal storage diseases are a heterogenous group of inherited disorders characterized by accumulation of undegraded substrates within lysosomes. Typically, these disorders are caused by an inability to perform an enzymatic reaction, often due to deficiency in the production or function of a lysosomal hydrolase. Sanfilippo (types A-D) is collectively one of the most common lysosomal storage diseases, with incidence estimates ranging from 1 in 25,000 to 100,000.^1,2^ Approximately fifteen percent of Sanfilippo disease is type B (*1*). The lysosomal storage material in Sanfilippo syndromes is the glycosaminoglycan heparan sulfate. The disease is associated with progressive neurodegeneration, typically with onset in early childhood (*2*). In the clinic, soluble lysosomal hydrolases are available in recombinant form for some types of lysosomal storage disease, and may be delivered by repeated infusion into the bloodstream or into the cerebrospinal fluid (*3-9*). However, no therapeutic options are currently approved for Sanfilippo syndrome.

To function as a therapeutic enzyme, a recombinant lysosomal hydrolase requires appropriate post-translational modification including glycosylation with mannose 6-phosphate (M6P) (*10*). Mannose 6-phosphate residues bind to mannose 6-phosphate receptors (M6PRs) expressed on the endosomal-lysosomal membrane, and results in transport of the enzyme from the Golgi apparatus to the lysosome, where it is active and catabolizes substrate in the acidic environment of this organelle. M6PRs are also present on the plasma membrane and mediate the endocytosis and subsequent trafficking of properly modified lysosomal enzymes from the extracellular milieu to the lysosome. One of the lysosomal hydrolases, the enzyme alpha-*N*-acetyl-glucosaminidase (NAGLU) that is responsible for Sanfilippo B syndrome (or mucopolysaccharidosis type IIIB, MPS IIIB), is not properly processed when produced recombinantly (*11, 12*). A fusion protein that contains the therapeutic enzyme (recombinant human NAGLU; rhNAGLU) plus a peptide fragment of IGF2, which can bind the cation-independent M6PR, forming rhNAGLU-IGF2 demonstrated improved intracellular uptake and reduction of lysosomal storage in Sanfilippo B fibroblasts (*13*). Using a mouse model, intracerebroventricular (ICV) delivery of rhNAGLU-IGF2 demonstrated therapeutic restoration of NAGLU, normalization of lysosomal storage, and dramatic improvement in markers of neurodegeneration and glial activation (*14*). Neonatal administration of rhNAGLU-IGF2 showed a treatment effect that lasted more than four weeks after a single ICV dose (*15*). A commercial form of rhNAGLU-IGF2 has reached clinical trials (*16-18*).

Recombinant enzymes delivered into cerebrospinal fluid can penetrate into deep brain structures and improve neuropathology in preclinical studies (*19-22*), and there is one recombinant therapeutic enzyme approved for intracerebroventricular administration in patients for the late-infantile form of neuronal ceroid lipofuscinosis (CLN2 disease) (*8*). Here, we studied the effect of intracerebroventricular rhNAGLU-IGF2 on behavior and survival in Sanfilippo B mice and studied effects upon biochemistry and proteomic profile of the brain of neonatal Sanfilippo B dogs.

## Methods

### Mouse model

*Naglu*-/- mice (B6.129S6-*Naglu*^*tm1Efn/J*^, backbred onto the C57BL6/J strain for at least 10 generations) were produced by crossing -/- male mice with +/- female mice, thus generating approximately 50% affected progeny. Mice received research grade rhNAGLU-IGF2 (100 µg), rhNAGLU (100 µg), or an equivalent volume (~5 µl) of vehicle (artificial cerebrospinal fluid (CSF)) in the left lateral ventricle beginning at PND 2 by manual injection. At four weeks of age, mice were cannulated into the left ventricle, and the infusion of enzyme/vehicle was given via cannulae at 28-day intervals until 16 weeks of age. To prevent hypersensitivity reactions, mice received premedication with diphenhydramine 5 mg/kg intraperitoneally 15 min prior to infusions. Research grade rhNAGLU-IGF2, rhNAGLU, and vehicle were provided by BioMarin Pharmaceutical, Inc. (Novato, CA). Experimental personnel were blinded as to treatment of either therapeutic enzymes or vehicle (coded “1, 2 or 3”). All animal experiments were approved by the institutional animal use committees at the Los Angeles Biomedical Research Institute at Harbor-UCLA Medical Center (now the Lundquist Institute) in Torrance, CA, and Washington University School of Medicine in St. Louis, MO.

### Auditory-evoked brainstem response (ABR) and electroretinography (ERG)

ABR were performed as described (*23*) using Tucker-Davis Technologies System 3 Complete ABR/OAE Workstation equipped with a flashlamp system and Medusa RA16 preamplifier and headstage, and a Hewlett-Packard PC computer running SigGen and BioSig software (Tucker-Davis Technologies, Alachua FL). Stimuli at each frequency and level were presented 1,000 times at 20/sec. The minimum sound pressure level required for detection of a response was determined using a 5 dB minimum step size. Flash ERG was performed as described previously (*23-25*). Flash ERG measurements were recorded on a Tucker-Davis System 3 Complete ABR/OAE Workstation (Tucker-Davis Technologies, Alachua, FL, USA) and analyzed using BioSig software. Mice were dark-adapted for 2h to obtain a mixed rod/cone ERG, determined from the average response to 5–10 presentations of light at 0.1 Hz. Mice were light-adapted for 20 min to obtain a pure cone ERG, obtained by measuring the average response following 50 light flashes at a frequency of 1 Hz. Comparisons were made of the b-wave amplitudes in nanovolts from the best of three measurements of each of the light-adapted and dark-adapted responses.

### Behavior

Male Sanfilippo B “mutant” mice and male carrier” mice were used for behavioral assessment at the UCLA Behavior Core. Elevated plus maze, novel object recognition, and social interaction tests were performed at the UCLA Behavioral Testing Core as previously described (*26*). The final infusion dose was given at 16 weeks of age, then infusion cannulae were removed before behavioral testing at 16-20 weeks of age was performed. Mice in these experiments were male and the implantation of cannulae prevented them from being caged with other males. To prevent confounding of behavioral experiments due to social isolation, these males were cohoused with a female companion and permitted to breed. This cohort of mice was euthanized at 20 weeks old following behavioral assessments.

Cohorts of mice of both sexes were studied at Washington University School of Medicine in St. Louis, Missouri. Activity in an open field was assessed at 22 weeks of age over 24 h in a rectangular polystyrene enclosure surrounded by photobeams monitored by computer software (MotorMonitor, Kinder Scientific, Poway, CA) (*27*). Testing consisted first of a 12 h block with the room lights off (“dark”) and ended with a 12 h block with the room lights on (“light”). Mice had access to food and water *ad libitum* during testing. Outcome measures included number of rearing events, distance traveled, and time spent resting. To test motor coordination (vestibular and cerebellar function), animals were tested on a rotarod (UGO Basile, Verese, Italy) at 32 and 40 weeks as described (*23*). Briefly, a rocking paradigm with reversal of direction of rotation after each full turn at 10 rpm for 3 minutes was executed for 3 consecutive days at 32 and 40 weeks of age. For each time point, training was provided for 2 consecutive days with 3 trials each day and 20 minutes of rest between each trial. Testing occurred on the third consecutive day and the length of time each mouse remained on the rod was recorded for all three trials and averaged. The mice were euthanized at 40 weeks old following behavioral assessments.

### Survival

In the survival experiment mutant mice were given monthly infusion starting at PND2 with vehicle (8 males, 8 females) or rhNAGLU-IGF2 (8 males, 8 females) as well as carrier mice given vehicle (3 males, 13 females) or rhNAGLU-IGF2 (8 males, 8 females). The survival experiment was carried to humane endpoints and terminated with the death of the last treated mutant mice.

### Biochemical assays

A subset of mice were used for biochemical analysis. Brains, liver, and heart were removed. Spinal cords were removed by ejection. Tissues were frozen on dry ice and stored at -80 °C until assay. NAGLU activity and total hexosaminidase activity were quantified using 4-methylumbelliferyl assays as previously described (*15*). Briefly the enzymatic activity of NAGLU and NAGLU–IGF2 was determined by hydrolysis of the fluorogenic substrate, 4-methylumbelliferyl-*N*-acetyl-α-glucosaminide, obtained from CalBiochem (now EMD Millipore Chemicals) or Toronto Research Chemicals, with minor modifications of a published protocol, using 1.0 mM substrate in the incubation mixture (*28*). Enzymatic activity of beta-hexosaminidase (combined A and B isoforms) was determined by hydrolysis of 4-methylumbelliferyl-*N*-acetyl-β-glucosaminide (EMD Millipore Chemicals, Darmstadt, Germany) using 1.25 mM substrate in the incubation mixture. For both enzymes, a unit of activity is defined as release of 1 nmol of 4-methylumbelliferone (4MU) per hour. Protein concentration was estimated by the Bradford method, using bovine serum albumin as a standard.

Analysis of heparan sulfate was performed by BioMarin Pharmaceutical (mouse tissues) or the UC San Diego GlycoAnalytics Core (canine tissues) as described previously (*29*). Tissues were homogenized and digested overnight with Pronase (Sigma-Aldrich, St. Louis, MO) in phosphate-buffered at 37 °C and centrifuged at 14,000 rpm for 20 min. The supernatant was passed through a DEAE column, and the bound GAG was eluted with 2 M NaCl. The GAG was then desalted on PD10-size exclusion column, and purified GAG was lyophilized. Dried GAG disaccharides were dissolved in aniline and reacted with freshly prepared sodium cyanoborohydride solution in a dimethyl sulfoxide/acetic acid mixture (7:3, v/v). Reactions were carried out at 65 °C for 1 h followed by 37 °C for 16 h. The samples were then dried using a speed vacuum at room temperature. Aniline-tagged disaccharides were separated on a C18 column using an ion pairing solvent mixture and analyzed by mass spectrometry in negative ion mode.

### Immunostaining

Mice were anesthetized with isoflurane and perfused transcardially with heparinized PBS followed by 4% paraformaldehyde in neutral PBS. After post-fixation in this solution for an additional 48 hours, the brains were removed and cryoprotected in 30% sucrose in 50 mM tris buffered saline, pH=7.6. Spinal cords were divided into cervical, thoracic and lumbosacral blocks, and 40µm coronal sections were cut throughout the cervical block on a Thermofisher HM430 freezing microtome (Thermofisher). A one in six series of 40 µm coronal brain sections from each mouse and a one in forty-eight series of spinal cord sections from each mouse were stained on Superfrost Plus slides using a modified immunofluorescence protocol for the following antibodies: rat anti-mouse CD68, 1:400 (Bio-Rad, Hercules, CA, cat # MCA1957), rabbit anti-LAMP1, 1:400 (Abcam, Cambridge, MA, cat# Ab24170). The intensity of immunostaining was assessed using a semi-automated thresholding image analysis method, as described previously, using Image Pro-Premier software (MediaCybernetics, Chicago, IL) (*30*). Unbiased optical fractionator counts of the number of Nissl-stained ventral horn neurons from the cervical spine were obtained as described previously (*30, 31*), using StereoInvestigator software (Microbrightfield, MBF Bioscience, Williston, VT) and a design-based optical fractionator method. A one in forty-eight series of 40µm coronal spinal cord sections from each mouse were stained with cresyl fast violet (Brand, City). Cells were sampled with counting frames (70 × 40 μm) distributed over a sampling grid of 150 × 150 μm superimposed over the region of interest at 100 × magnification. Some vehicle-treated mutant and carrier control samples had undergone ex vivo imaging and were therefore not suitable for Nissl staining, so age-matched, untreated, mutant and carrier control mice were used.

### Retinal histology

Eyes were obtained mice and immersion fixed in 2% glutaraldehyde/4% formaldehyde in PBS. Samples were embedded in Epon-Araldite resin and sectioned into 1-micron-thick sections and stained with toluidine blue as described (*23*). After careful alignment of orientation, the retinal thickness and the thickness of each layer were measured using ImageJ (*32*). Each measurement (e.g., length of outer nuclear layer, length of rods and cones layer, etc.) was performed in triplicate in ten locations along the axis of the eye for each mouse and the results averaged for each mouse.

### Canine model

This work was performed in accordance with the *Guide for the Care and Use of Laboratory Animals* and approved by the Iowa State University institutional Animal Care and Use Committee (IACUC). Affected dogs were anesthetized and a 22-gauge needle inserted into the cisterna magna. CSF (0.2 to 0.5 ml) was collected free-flowing. A syringe was attached to the needle hub to deliver rhNAGLU-IGF2 at 1 week of age and again at 4 weeks of age (in sterile saline, total volume 300 μl at 1 week and 700 μl at 4 weeks). Dogs were euthanized at 8 weeks of age (4 weeks after the second dose). Dose was calculated per estimated brain weight. At one week of age, canine brains are roughly 1/38 of body weight, and the administered dose was 400 mg rhNAGLU-IGF2/kg brain, or approximately 7 mg. For a 2.5 kg four week-old pup, in which brain weight is 1/60 of body weight, the dose was approximately 17 mg. Control group animals were not treated. At necropsy, terminal CSF and blood were collected, and dogs were perfused with chilled PBS and brains removed. Brains were sliced coronally into 4 mm slabs using a pre-chilled, stainless steel canine brain matrix (ASI Instruments) as described elsewhere (*33*). A 3 mm biopsy punch was used to collect samples from cerebral gray matter, cerebral white matter, caudate, and cerebellar vermis. Samples were snap-frozen in an isopentane bath on dry ice.

### Proteomics

Snap-frozen canine samples (approximately 1×1×0.4 cm) from a parasagittal gyrus, including gray matter and subcortical white matter were analyzed by LC/MS/MS at Caltech. Protein lysates from canine brain cortex samples were prepared using Thermo EasyPrep Mini MS Sample Prep Kit (cat# A4006). Protein concentrations were determined using Bradford (Bio-Rad) and 100 μg total protein from each sample were prepared for mass spectrometry acquisition using Thermo EasyPep Mini MS Sample Prep Kit. After determining peptide concentrations by Pierce Quantitative Fluorometric Peptide Assay (cat# 23290), 20 μg peptide from each sample was labeled with TMT10plex Isobaric Label Reagent Set (ThermoFisher Scientific, cat# A37725) following to the manufacture’ s instruction. Labeled samples were combined and dried using vacuum centrifugation. Samples were then separated into 8 fractions using the High pH reversed-phase peptide Fractionation Kit (ThermoFisher Scientific, cat# 84868). The fractions were dissolved with 0.1% formic acid and peptide concentration was determined with Quantitative Colorimetric Peptide Assay (ThermoFisher Scientific, cat# 23275).

TMT labeling LC-MS/MS experiments were performed using an EASY-nLC 1000 connected to an Orbitrap Eclipse Tribrid mass spectrometer (Thermo Scientific). 0.5 μg of each fraction was loaded onto an Aurora AUR2-25075C18A UHPLC Column (IonOpticks, Fitzroy, Australia) and separated over 131 min at a flow rate of 0.4 μL/min with the following gradient: 2-6% Solvent B (2 min), 6-22% B (78 min), 22-50% B (40 min), 50-95% B (1 min), and 95% B (10 min). Solvent A consisted of 97.8% H_2_O, 2% ACN, and 0.2% formic acid, and solvent B consisted of 19.8% H_2_O, 80% ACN, and 0.2% formic acid. An MS1 scan was acquired in the Orbitrap at 120k resolution with a scan range of 400-1600 m/z. The AGC target was 1 × 10^6^, and the maximum injection time was 50 ms. Dynamic exclusion was set to exclude features after 1 time for 60 s with a 10-ppm mass tolerance. MS2 scans were acquired with CID activation type with the IonTrap. The isolation window was 0.7 m/z, the collision energy was 35%, the maximum injection time was 35 ms and the AGC target was 1 × 10^4^. MS3 scans were acquired with HCD activation type in the Orbitrap at 60k resolution with a scan range of 100-500 m/z. The isolation window was 0.7 m/z, the collision energy was 65%, the maximum injection time was 118 ms and the AGC target was 2.5 × 10^5^. Ion source settings were as follows: ion source type, NSI; spray voltage, 2300 V; ion transfer tube temperature, 300°C. System control and data collection were performed by Xcalibur software (ThermoFisher Scientific).

### Data analysis

Graphs were produced on GraphPad Prism (version 9 or 9.1; GraphPad Software, San Diego, CA) or SigmaPlot version 14.0 (Systat Software Inc., San Jose, CA). Linear regression was performed using SigmaPlot. Statistical analysis of continuous variables was performed on Stata (version 16 or 17) or GraphPad Prism using ANOVA (if normal distribution), or if non-normal using Mann-Whitney. The proteomic data processing was performed through Proteome Discoverer 2.4 (ThermoFisher Scientific) using Uniprot canine database and the SequestHT with Percolator validation (*34, 35*). Normalization was performed relative to the total peptide amount. Further analyses were performed as below: limma analyses were performed using R studio following the user guide (*36*); volcano plot was generated with Origin 2019b; DE proteins were tested for enrichment analysis using g:Profiler (*37*) and plotted with Prism 7 (GraphPad Software, San Diego, CA); heatmaps were generated with Prism 7.

## Results

### Lysosomal storage

Male and female mice received ICV injections of rhNAGLU-IGF2, rhNAGLU, or vehicle at PND2. They were cannulated at 4 weeks of age and received therapeutic enzyme or vehicle via cannulae at monthly intervals until 16 weeks of age and euthanized 4 weeks later. Brain and heart showed a reduction in HS that reached statistical significance for each treatment group compared to vehicle-treated Sanfilippo B mice (p<0.001, **Fig. 1**). All intergroup differences were statistically significant by ANOVA. There was some residual NAGLU enzymatic activity detected in mouse brains in both treatment groups despite the 28-day interval from the final dose and tissue collection (**Supplemental Table S1**). Similar to previous studies, we found a reduction in the disease-associated marker beta-hexosaminidase in brain, liver, heart, and spinal cord of mice treated with rhNAGLU (p=0.0016) or rhNAGLU-IGF2 (p<0.001) compared to vehicle-treated Sanfilippo B mice (**Table 1**).

**Table 1.** Beta-Hexosaminidase Activity in Mice (28 days post-dose)

|  | Brain<br><u>1+2</u> | Brain<br><u>3</u> | Brain<br><u>4</u> | Brain<br><u>5+6</u> | Spinal<br><u>Cord</u> | <u>Liver</u> | <u>Heart</u> |
| --- | --- | --- | --- | --- | --- | --- | --- |
| Mutant +<br>Vehicle | 3687 ±<br>244 | 3944 ±<br>280 | 3305 ±<br>112 | 2258 ±<br>276 | 2058 ±<br>273 | 3377 ±<br>907 | 1847 ±<br>296 |
| Carrier +<br>Vehicle | 1318 ±<br>136 | 1420 ±<br>144 | 1307 ±<br>180 | 874 ±<br>220 | 746 ±<br>59.8 | 435 ±<br>35.6 | 190 ±<br>64.7 |
| Mutant +<br>NAGLU | 1681 ±<br>173 | 1681 ±<br>67.6 | 1708 ±<br>106 | 955 ±<br>244 | 701 ±<br>74.3 | 755 ±<br>133 | 721 ±<br>335 |
| Mutant +<br>NAG-IGF2 | 2052 ±<br>265 | 1884 ±<br>321 | 1820 ±<br>320 | 1076 ±<br>258 | 706 ±<br>80.3 | 980 ±<br>307 | 858 ±<br>208 |

Values are means ± standard deviations and are expressed as units of activity per milligram protein in homogenized tissue. Brains were divided coronally and numbered rostral to caudal 1- 6, with the 1<sup>st</sup> and 2<sup>nd</sup> combined and the 5<sup>th</sup> and 6<sup>th</sup> combined for the assay. NAG-IGF2: rhNAGLU-IGF2, NAGLU: rhNAGLU. All values in the rhNAGLU or rhNAGLU-IGF2-treated mutant mice were statistically significant compared with vehicle-treated mutant mice by Mann- Whitney.

**Fig. 1.**
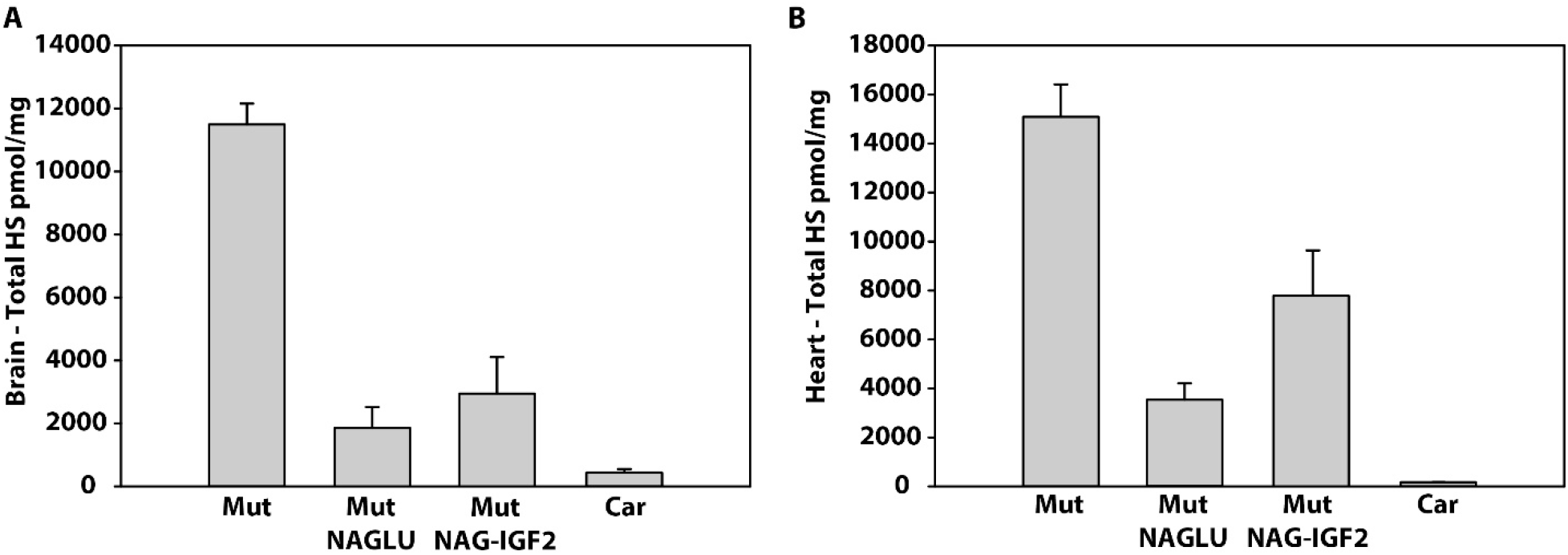
Heparan sulfate (HS) concentrations in the **(A)** brain and **(B)** heart of experimental mice and controls. Mut: Sanfilippo B (*Naglu-/-*) mice. Car: heterozygous carrier (*Naglu +/-*) mice that are expected to be phenotypically unaffected. NAG-IGF2: rhNAGLU-IGF2, NAGLU: rhNAGLU. Mutant and Carrier control mice were treated with vehicle, and treatments and vehicle were administered intracerebroventricularly as described in the methods. Units are expressed in mg of wet weight, with means and standard deviation shown. N=8 per group. Uncorrected p-value 0.0002 for Mut vs. Mut NAGLU or Mut NAG-IGF2 in both brain and heart by Mann-Whitney.

We also found decreased immunoreactivity for CD68, a marker of activated microglia, and LAMP1, a marker of lysosomes, in the primary somatosensory cortex (S1BF) and striatum (caudate and putamen) of mice treated with rhNAGLU or rhNAGLU-IGF2 (**Figs. 2 and 3**). Carrier mice treated with rhNAGLU or rhNAGLU-IGF2 did not differ from vehicle-treated carrier mice in LAMP1 or CD68 immunostaining in either brain region (not shown).

**Fig. 2.**
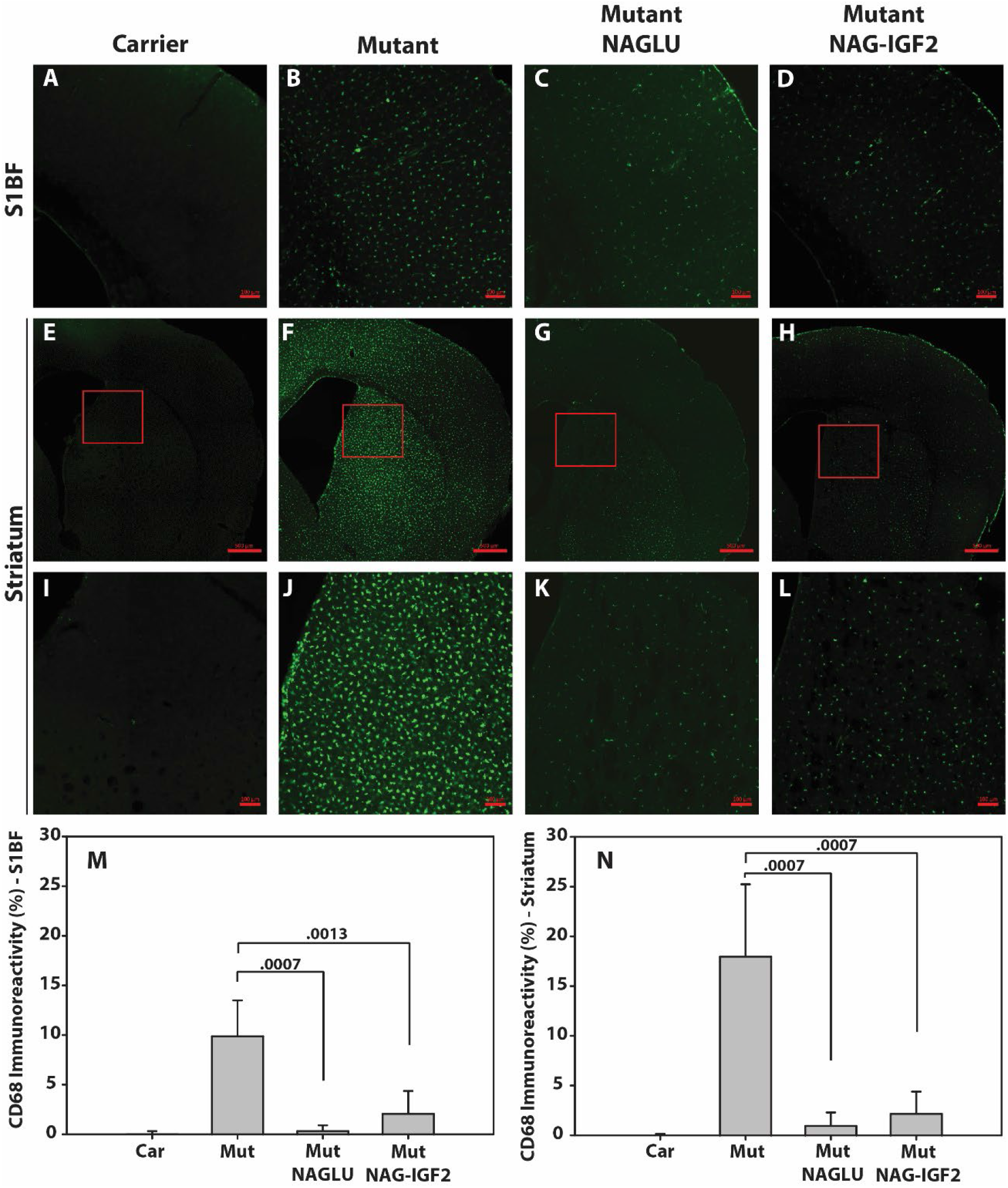
CD68 immunoreactivity in mouse brain. **(A-L)** Immunofluorescence staining with anti-CD68 (green) in the primary somatosensory cortex (S1BF) and striatum (caudate and putamen) in mice. Red boxes indicate insets for the panels below. Scalebar 100 µm in panels A-D and I-L. Scalebar 500 µm in panels E-H. (**M-N**) Bar graphs depict means and standard deviations of thresholding image analysis. N=4-9 per group. Uncorrected p-values show between-group comparisons by Mann-Whitney. Mut (mutant): Sanfilippo B (*Naglu-/-*) mice. Car (carrier): heterozygous (*Naglu +/-*) mice. NAGLU: rhNAGLU. NAG-IGF2: rhNAGLU-IGF2. Mutant and Carrier control mice were treated with vehicle as described in the methods.

**Fig. 3.**
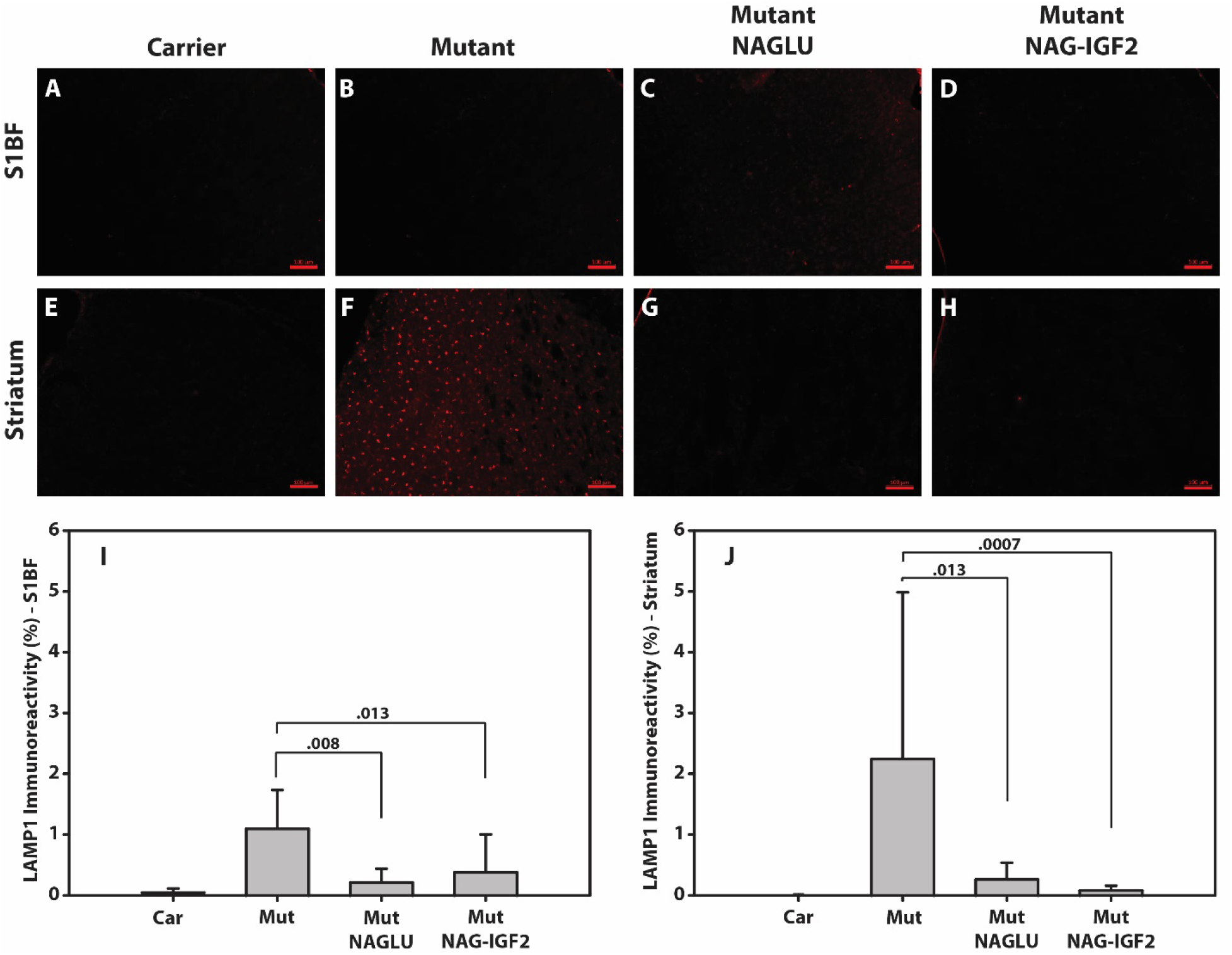
LAMP1 immunoreactivity in mouse brain. **(A-H)** Immunofluorescence staining with anti-LAMP1 (red) in the primary somatosensory cortex (S1BF) and striatum (caudate and putamen) in mice. Scalebar 100 µm. **(I-J)** Bar graphs depict means and standard deviations of thresholding image analysis. N=4-9 per group. Uncorrected p-values show between-group comparisons by Mann-Whitney. Mut (mutant): Sanfilippo B (*Naglu-/-*) mice. Car (carrier): heterozygous (*Naglu +/-*) mice. NAGLU: rhNAGLU. NAG-IGF2: rhNAGLU-IGF2. Mutant and Carrier control mice were treated with vehicle as described in the methods.

### Spinal cord

Neuron counts of cervical spinal ventral horn were performed in a subset of mice. Sanfilippo B mice showed 18% lower mean estimated neuron counts compared to carrier controls (p=0.021, **Fig. 4**). Mice treated with rhNAGLU or rhNAGLU-IGF2 showed mean neuron counts that were not significantly different from carrier mice. The treated mice in both groups showed higher spinal neuron counts compared to untreated Sanfilippo B mice, and this reached statistical significance for mice treated with rhNAGLU (p=0.014) but not rhNAGLU-IGF2 (p=0.088).

**Fig. 4.**
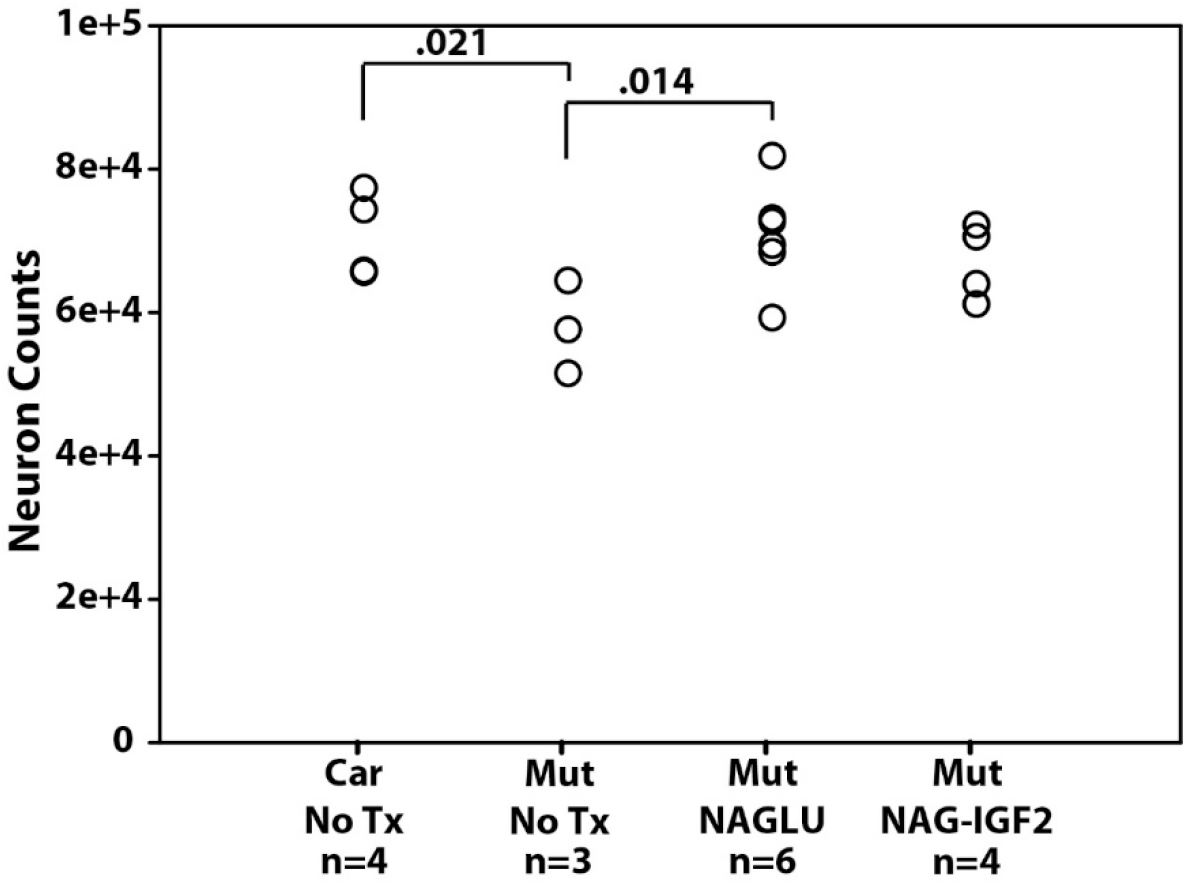
Neuron counts in mouse cervical spinal cord. Circles represent individual mice. Mut (mutant): Sanfilippo B (*Naglu-/-*) mice. Car (carrier): heterozygous (*Naglu +/-*) mice. NAGLU: rhNAGLU. NAG-IGF2: rhNAGLU-IGF2. Mutant and Carrier control mice were not treated (“No Tx”). Unadjusted p-values show pairwise comparisons by ANOVA.

Sanfilippo B mice showed increased CD68 staining compared to normal controls suggestive of activated microglia in cervical, thoracic, and lumbar gray matter of the spinal cord (**Supplemental Figure S1**). CD68 staining for activated macrophages in cervical spinal gray matter of Sanfilippo B mice showed consistent increases in the spinal cord dorsal and ventral horns compared to heterozygous normal mice or mice treated with rhNAGLU or rhNAGLU-IGF2 (**Fig. 5**). Similarly, LAMP1 immunostaining of the cervical spinal gray matter of Sanfilippo B mice showed increased staining qualitatively and quantitatively by threshold analysis of immunofluorescence signal in dorsal and ventral horns of the spinal cord compared to unaffected carrier mice (**Fig. 5**). Quantification of both markers in mice treated with rhNAGLU or rhNAGLU-IGF2 showed levels of immunostaining that were similar to those of unaffected carrier mice, with no significant differences in immunoreactivity between groups of mice treated with rhNAGLU and rhNAGLU-IGF2. Carrier mice treated with rhNAGLU or rhNAGLU-IGF2 did not differ from vehicle-treated carrier mice in LAMP1 or CD68 immunostaining (not shown).

**Fig. 5.**
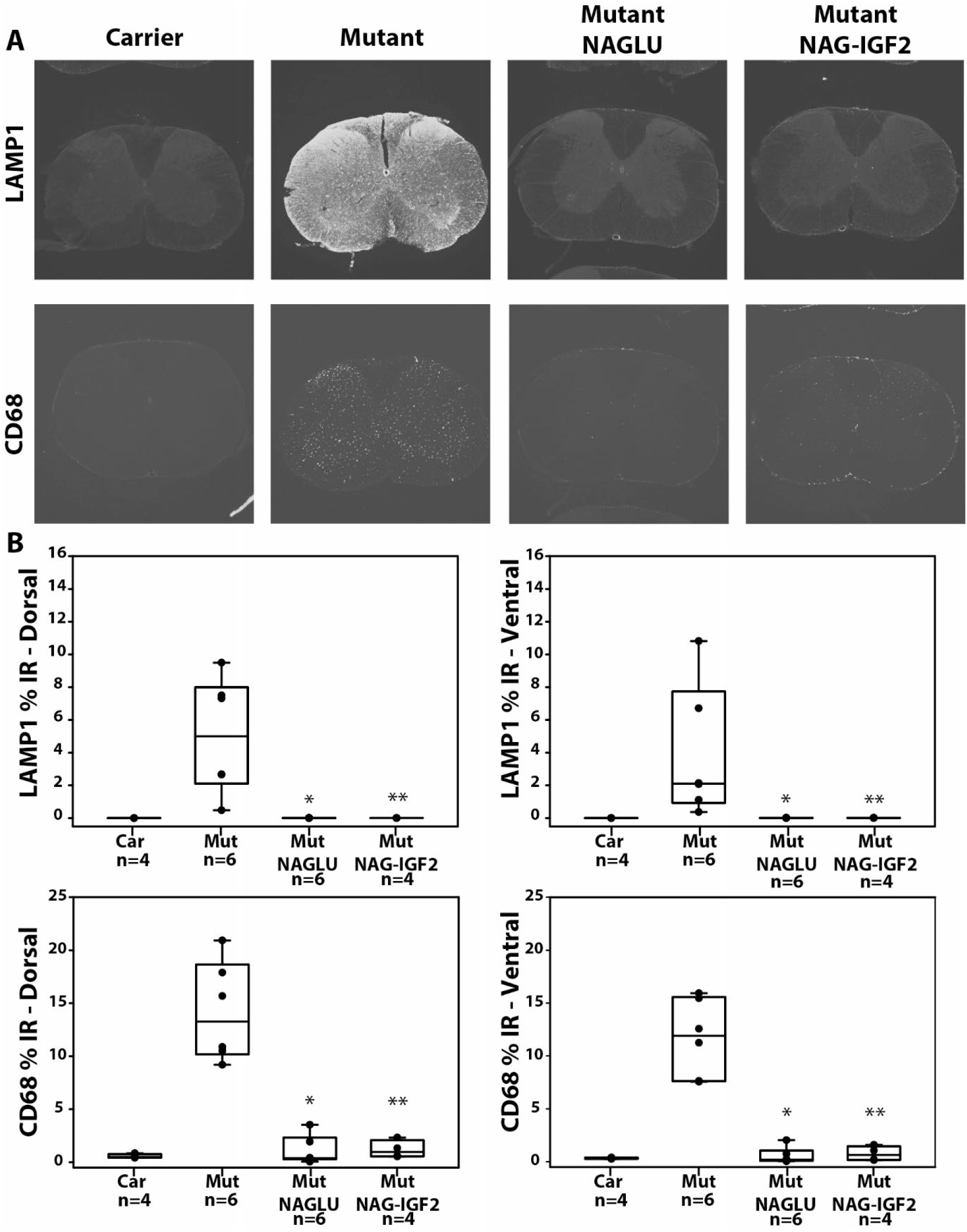
CD68 and LAMP1 immunostaining in mouse cervical spinal cord. **A**) Representative immunofluorescence images of axial sections of spinal cord stained for LAMP1 and CD68 (10x magnification). **B**) Box plots showing median, first and third quartiles, and range of immunoreactivity (IR) in dorsal (upper panels) and ventral (lower panels) cervical cord. Dots represent individual mice. Mut (mutant): Sanfilippo B (*Naglu-/-*) mice. Car (carrier): heterozygous (*Naglu +/-*) mice. NAGLU: rhNAGLU. NAG-IGF2: rhNAGLU-IGF2. Mutant and Carrier control mice were treated with vehicle as described in the methods. *p=0.0022 and **p=0.0095 vs. Mut by Mann-Whitney (uncorrected).

### Behavior

Male Sanfilippo B and control mice were treated with rhNAGLU, rhNAGLU-IGF2, or vehicle from PND2 to 16 weeks as described in the methods. We previously found that Sanfilippo B mice show reduced measures associated with fear in mice, as evidenced by decreased stretch attend postures on the elevated plus maze (*26*). This difference between normal and Sanfilippo B mice was again seen here (**Fig. 4**), with a more normal number of stretch attend postures in Sanfilippo B mice treated with either rhNAGLU or rhNAGLU-IGF2.

We did not find significant intergroup differences between vehicle-treated Sanfilippo B mice and controls in radial arm maze performance, social interaction testing, dark/light activity, or novel object recognition testing, such that these measures were not informative in assessing a therapeutic response (**Supplemental Figures S2-S6**).

### Survival

Sanfilippo B and control mice were treated with rhNAGLU-IGF2 or vehicle (four groups; **Fig. 6**). The experiment was terminated after the last vehicle-treated mutant mouse died. Results indicate a 58% increase in survival in Sanfilippo B mice treated with rhNAGLU-IGF2 (mean 52w, median 48w) compared to vehicle-treated Sanfilippo B mice (mean 33w, median 37w; p<0.001). Causes of death and other adverse events (mainly loss of cannulae and reimplantation) are described in **Supplemental Table S3**.

**Fig. 6.**
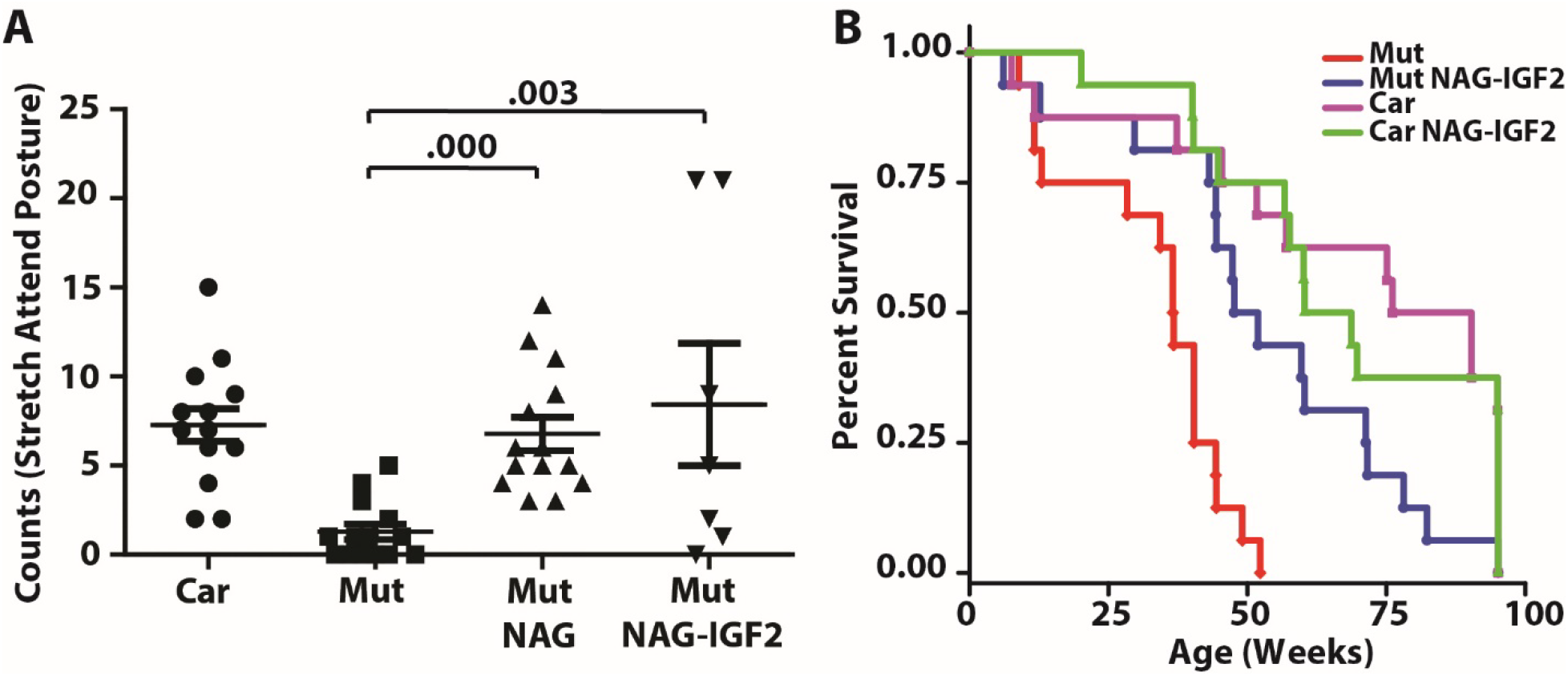
Fear and Survival. A) Count of stretch attend postures during testing on the elevated plus maze. Mut (mutant): Sanfilippo B (*Naglu-/-*) mice. Car (carrier): heterozygous (*Naglu +/-*) mice. NAGLU: rhNAGLU. NAG-IGF2: rhNAGLU-IGF2. Mutant and Carrier control mice were treated with vehicle as described in the methods. Carrier mice treated with rhNAGLU or rhNAGLU-IGF2 did not differ from carrier mice treated with vehicle (not shown). Each symbol = one mouse, with means and s.e.m. shown; N=7-14 per group. Numbers above brackets indicate uncorrected p-values by Mann-Whitney. B) Survival curves for Sanfilippo B mice (Mut) and carrier controls (Car) treated with ICV vehicle (Veh) or rhNAGLU-IGF2 (NAG-IGF2) monthly from PND 2. N=16 per group, males and females. p<0.001 Mut-Veh versus Mut-NAG-IGF2.

### Motor Coordination

Rotarod testing was performed in these mice at age 32 and 40 weeks, and results are shown in **Fig. 7**. Sanfilippo B mice treated with vehicle showed slightly reduced rotarod performance at 32 weeks (p=0.0026), with further worsening at 40 weeks (p=0.004). Sanfilippo B mice treated with rhNAGLU-IGF2 showed improved rotarod performance compared to vehicle-treated Sanfilippo B mice. Performance of the enzyme-treated mice was similar to normal mice at age 32 weeks. At 40 weeks, treated mice performed significantly better than vehicle-treated Sanfilippo B mice (p=0.002), but were also significantly different from unaffected carrier controls (p=0.006). One Sanfilippo B mouse treated with vehicle and one carrier mouse treated with rhNAGLU-IGF2 died between weeks 32 and 40.

**Fig. 7.**
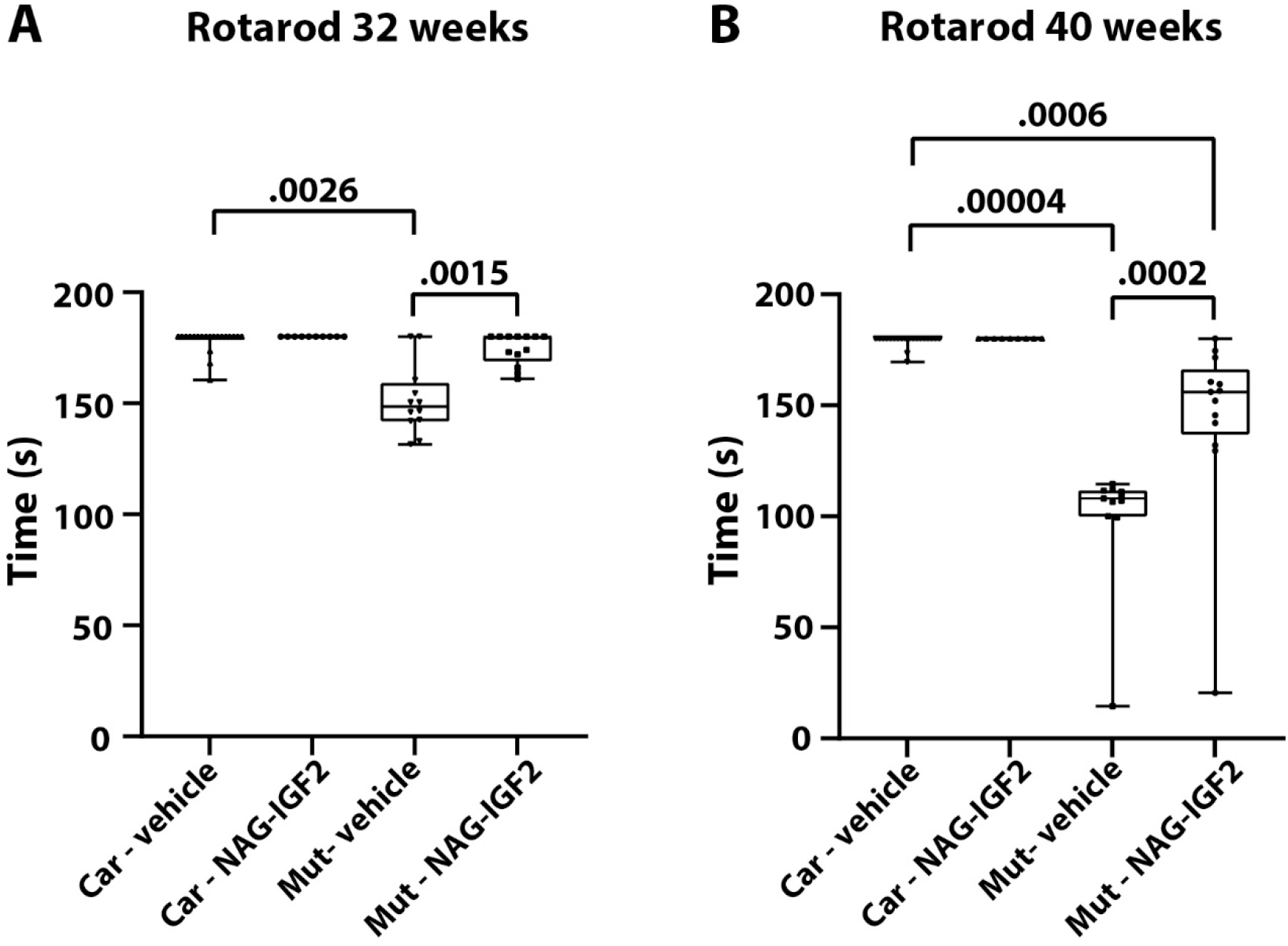
Time to fall off rotarod in Carrier (“Car,” *Naglu+/-*) and Mutant (“Mut,” *Naglu*-/-) mice treated with vehicle or rhNAGLU-IGF2 (NAG-IGF2) ICV monthly from PND 2. Mice were studied at (**A**) 32 and (**B**) 40 weeks. Mut – Vehicle, n=12 at 32 weeks and n=11 at 40 weeks; Mut – NAG-IGF2, n=13; Car – vehicle, n=20; Carrier – NAG-IGF2, n=10 at 32 weeks and n=9 at 40 weeks. Dots show individual mice, boxes outline the median, 25^th^ and 75^th^ centile, and whiskers depict range. Numbers above brackets indicate p-values by Mann-Whitney.

### Hearing and vision

Acoustic brainstem responses did not show intergroup differences between Sanfilippo B mice and unaffected carrier mice (**Supplemental Figure S7**). Electroretinographs showed retinopathy in Sanfilippo B mice that persisted despite treatment with rhNAGLU-IGF2 (**Fig. 8**). Cross-sectional analysis of the thickness of different layers of the retina post-mortem showed reduced thickness in Sanfilippo B mice compared to carrier controls, which reached statistical significance for the inner plexiform layer, outer nuclear layer, and rods and cones (uncorrected p-values 0.040, 0.008, and 0.038, respectively). Mice treated with rhNAGLU-IGF2 showed increased thickness of the individual retinal layers, reaching statistical significance for the outer nuclear layer (uncorrected p value 0.035).

**Fig. 8.**
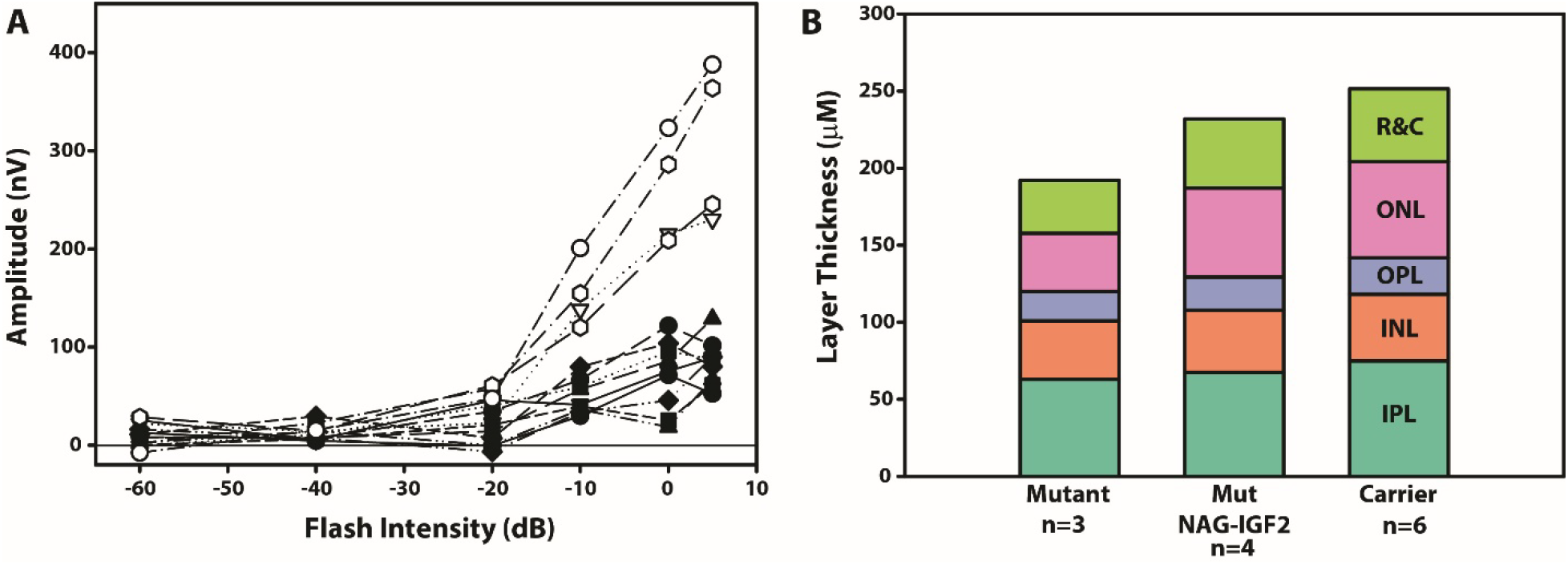
Retinal disease in Sanfilippo B mice. **A)** Electroretinogram (ERG) in Sanfilippo B mice treated with rhNAGLU-IGF2 (filled symbols) or carrier mice treated with vehicle (open symbols). **B**) Retinal thickness by layer in Sanfilippo B mice treated with rhNAGLU-IGF2 (NAG-IGF2). IPL: inner plexiform layer; INL: inner nuclear layer; OPL: outer plexiform layer; ONL: outer nuclear layer; R&C: rods and cones. Mutant: Sanfilippo B (*Naglu-/-*) mice. Carrier: heterozygous (*Naglu +/-*) mice. NAG-IGF2: rhNAGLU-IGF2. Mutant and Carrier control mice were treated with vehicle as described in the methods.

### Canine studies

This is naturally-occurring canine model of MPS IIIB, first identified in Schipperke dogs, which provides advantages of a larger, more complex brain in which to test therapeutic enzymes (*38, 39*). In this experiment, three Sanfilippo B dogs received two doses of rhNAGLU-IGF2 (~7 mg at 1 week and ~17 mg at 4 weeks of age) into the cerebrospinal fluid at the cisterna magna and were euthanized at 8 weeks of age, four weeks following the second and final dose. Three untreated Sanfilippo B dogs and three untreated carrier dogs were used as controls. Sanfilippo B dogs showed increased HS and beta-hexosaminidase levels in most brain regions compared to controls (**Fig. 9**). These disease-associated changes were reduced essentially to normal levels in treated animals, even in deeper regions of brain such as white matter and caudate nucleus. Some NAGLU activity was detected in brain tissue despite a four-week interval from the date of the final rhNAGLU-IGF2 dose and tissue collection (**Supplemental Table S3**).

**Fig. 9.**
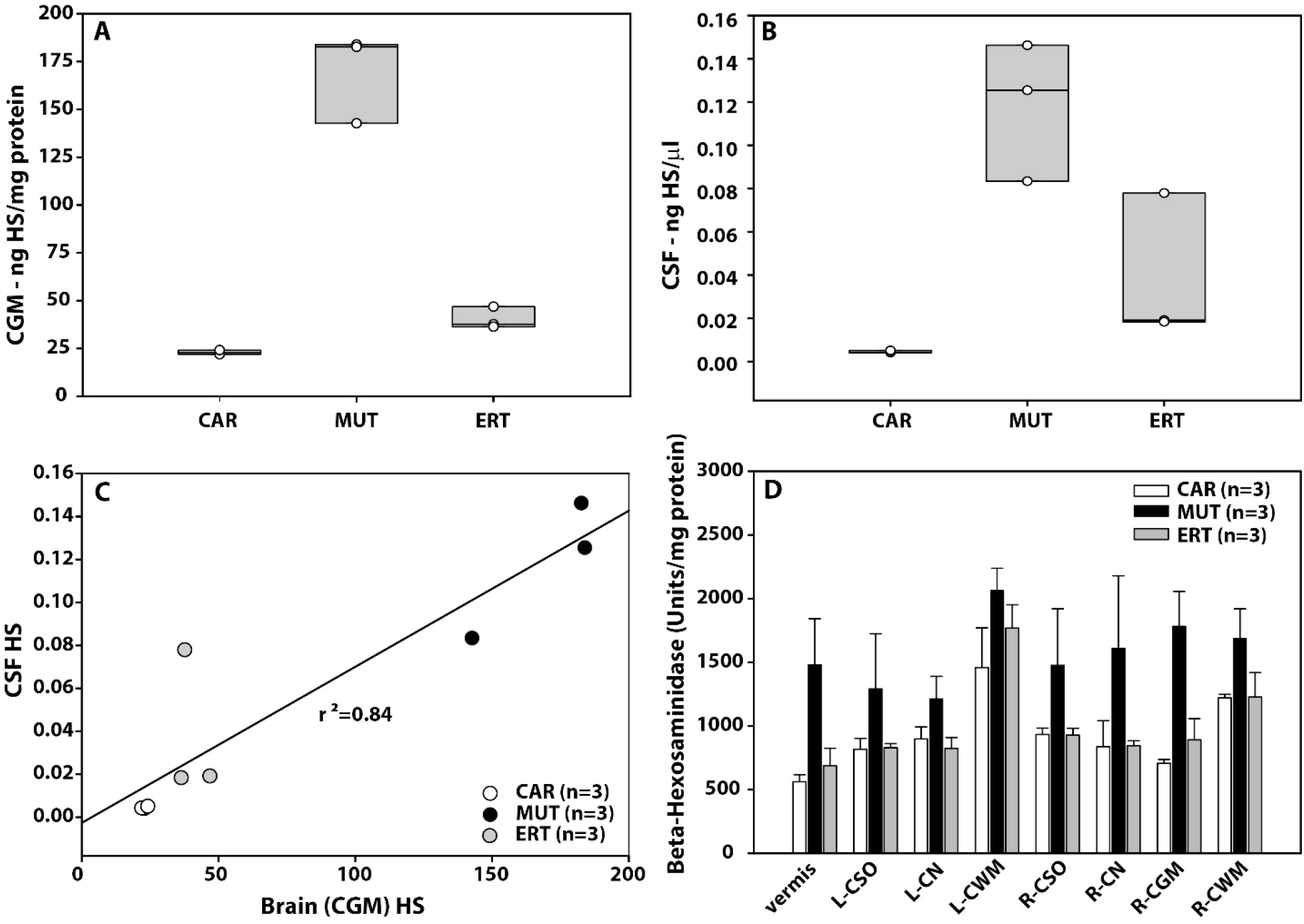
Canine Sanfilippo B. Sanfilippo B dogs (MUT) and carrier controls (CAR) were treated with intra-cisternal rhNAGLU-IGF2 (ERT) or untreated at 1 and 4 weeks of age and euthanized at 8 weeks (n=3 per group, both sexes). **(A)** Box plot of heparan sulfate (HS) in left cerebral gray matter (CGM). **(B)** Box plot of heparan sulfate (HS) in CSF. **(C)** Regression curve showing HS in cerebrospinal fluid (CSF) versus HS in CGM. Open circles represent individual animals. (D) Beta-hexosaminidase in brain regions (means and standard deviation). CSO: centrum semiovale. CN: caudate nucleus. CGM: cerebral gray matter. CWM: cerebral white matter.

Proteomic analysis was performed on cerebral cortical samples of each animal at the Chou lab as described in the Methods (**Fig. 10**). Tandem mass tag (TMT) 10-plexed labelling was employed to evaluate the proteomic changes in Sanfilippo B mutant animals. Specifically, we prepared three independent biological wild type (CAR), Sanfilippo B (MUT) and rhNAGLU-IGF2-treated Sanfilippo B (ERT) canine brain cortex samples. The TMT labeled samples were pooled and further fractionated into 8 fractions and analyzed with the RTS-SPS-MS3 method(*40*). A total of 5806 proteins were identified, and 4442 of these proteins were quantified across all 9 samples (**Supplemental Table S4**). We identified 309 proteins that demonstrated significantly different quantities (p < 0.05) between the untreated Sanfilippo B (MUT) and wild type (CAR) samples (**Supplemental Table S5**) by limma analysis. Among them, a set of 41 proteins with log_2_ (fold change) >0.5 (up-regulated) or <-0.5 (down-regulated) was classified as the final differentially expressed proteins (DEPs) (**Fig. 10A, Supplemental Table S6**). Subsequently, functional enrichment analysis was performed on these 41 DEPs. The top four relevant changed cellular components in the significantly enriched GO terms and top four changed KEGG or REAC functional pathways are presented in **Fig. 10B**. The GO: CC analysis showed these DEPs were mainly located at lysosome, secretory granule, extracellular matrix, and in vesicles. Pathway analysis indicated the DEPs primarily related to Glycosaminoglycan degradation, Keratan sulfate degradation, Neutrophil degranulation, and Innate Immune System. As shown in **Fig. 10C**, ERT therapy reversed several proteins that were down-regulated in mutant animals compared to carrier controls, such as DPP7 and CALB2. Notably, most of the elevated protein expression seen in Sanfilippo B canines were reversed by ERT, including GNS, LAMP1, HEXB, and CD63, among others. These results provide valuable insights for identifying novel hallmarks of lysosomal diseases and therapeutic targets for Sanfilippo B.

**Fig. 10.**
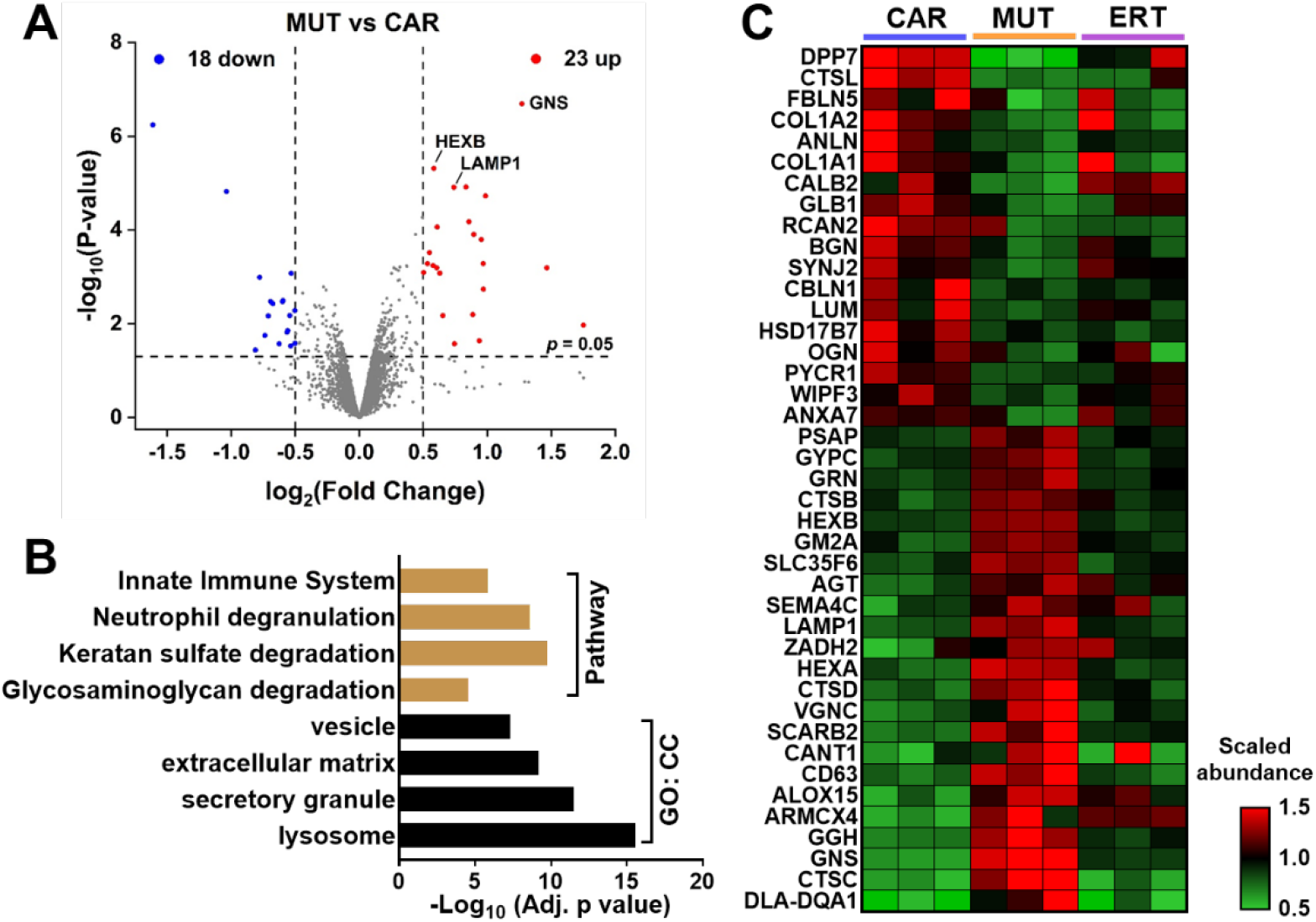
Proteomic analysis in canine Sanfilippo B. Experimental animals were the same as in Fig. 9 and included heterozygous carriers (CAR), Sanfilippo B (MUT), and rhNAGLU-IGF2-treated Sanfilippo B (ERT) groups. Proteins were analyzed from canine brain cortex. **(A)** Volcano plot displaying the proteomic changes in Sanfilippo B mutant, n=3. **(B)** Functional enrichment analysis of the differently expressed proteins affected by Sanfilippo B mutant. **(C)** Heatmap displaying the scaled abundance of the differently expressed proteins affected by Sanfilippo B mutant and rhNAGLU-IGF2 treatment.

## Discussion

Early treatment of lysosomal disease is necessary for maximum therapeutic effect. Here, we demonstrated that early treatment of Sanfililppo B mice and dogs with ICV enzyme replacement therapy corrects biochemical abnormalities, neuropathology, behavior, and survival. Our behavioral assessments were designed to separately evaluate the behaviors that might be impacted by physical disease, such as motor activity and balance, from behaviors that are primarily representative of altered central nervous system function, such as fear behavior. We found evidence that rhNAGLU-IGF2 administration improved survival, physical disease, and fear behavior in Sanfilippo B mice.

The IGF2 peptide was included to provide uptake of NAGLU via the IGF2-MPR, because rhNAGLU (unlike naturally-occurring NAGLU) lacks mannose 6-phosphate. Uptake and targeting via the MPR is an essential property of lysosomal enzymes. Neuronal cell lines cultured *in vitro* rely on M6PRs for intracellular uptake, suggesting that treating these disease-critical cells requires the use of this system (*15, 41, 42*). However, in this case we did not find a therapeutic advantage to ICV rhNAGLU-IGF2 compared to rhNAGLU, although direct comparison was limited in our study. A possible explanation for our findings is that the chosen dose and/or frequency did not reveal differences between the efficacy of rhNAGLU and rhNAGLU-IGF2 that might otherwise have been apparent. If the dose and frequency were higher than necessary, nonspecific enzyme uptake would be expected to occur and possibly eliminate the predicted differences between the native and fusion protein forms of enzyme. Alternatively, if the dose were too low, microglial and perivascular cell uptake could have been predominant in both the rhNAGLU and rhNAGLU-IGF2-treated groups. We previously demonstrated *in vivo* in a murine model of Sly syndrome (mucopolysaccharidosis type VII) that glycosaminoglycan levels and secondary enzyme elevations can be significantly reduced in the absence of any demonstrable cellular enzyme uptake ((*43*). *In vitro* and *in vivo* studies of Sanfilippo B and other lysosomal disorders have demonstrated that macrophage-lineage cells do not depend entirely on M6PRs to engulf material from the extracellular milieu (*44, 45*), and this has also been observed for mouse microglial cells incubated with rhNAGLU (*41*). In this scenario, both rhNAGLU and rhNAGLU-IGF2 would have treated mainly these glial cells and not sufficiently treated the neurons directly, resulting in a phenotypic improvement corresponding to correction in glial and other supportive cells only. Even with enzymes that are well mannose 6-phosphorylated, such as recombinant human alpha-L-iduronidase (for Hurler syndrome also known as MPS I) for example, distribution studies of intravenously administered enzyme showed preferential localization in sinusoidal cells with lesser staining appearing in hepatocytes. There could be a “threshold effect” at work, in which saturating macrophage-lineage cells is necessary prior to sufficient uptake into the target cells. It is not clear whether rhNAGLU or rhNAGLU-IGF2 were taken up into neurons *in vivo* in the brain during the experiments described here, because we were not able to assess biodistribution in our studies for the reasons described above. Previously published studies of rhNAGLU-IGF2 did show staining for NAGLU in neurons (*14*), but the dosing interval for that study was higher – four doses in two weeks – compared to the monthly dosing used here. If monthly dose and frequency of administration of rhNAGLU-IGF2 was inadequate to treat neurons yet sufficient to induce behavioral improvements, this implies that our phenotypic read-outs in Sanfilippo B mice are not sufficient to measure the neurological disease process and the impact of treatment upon this.

Besides enzyme replacement therapy, another approach in development is gene therapy to deliver the coding sequence for *NAGLU* as a theoretically permanent means of correction. Studies of adeno-associated viral vectors used to deliver NAGLU to Sanfilippo B mice show improvements in HS, neuropathology, behavior, and survival that are similar to those reported here for recombinant enzyme administration (*46, 47*). However, it is not known whether cells that are transduced by viral vectors bearing the *NAGLU* transgene produce NAGLU that is properly mannose 6-phosphorylated. Nevertheless, *in vitro* studies of NAGLU secreted by cells transduced using adeno-associated viral vector-2 demonstrated that the secreted NAGLU was able to correct intracellular HS accumulation as evidenced by an ^35^ SO_4_ assay (*48*). These authors did not study competitive inhibition of this effect by mannose 6-phosphate (*49*), so it is not clear whether the enzyme used that receptor system for uptake. It is likely that the concentration of enzyme that is required for correction of glycosaminoglycan accumulation is far lower than that required to demonstrate intracellular uptake of active enzyme (*50*). Both rhNAGLU-IGF2 and rhNAGLU were capable of reducing intracellular HS in human Sanfilippo B fibroblasts *in vitro*, although 10-fold higher concentrations of rhNAGLU compared to rhNAGLU-IGF2 were required to achieve this effect (*41*). To study mannose 6-phosphorylation of NAGLU in the context of gene therapy, induced pluripotent stem cells derived from Sanfilippo B mouse embryonic fibroblasts were transduced with lentivirus bearing the human *NAGLU* transgene under a cytomegalovirus promoter (*51*). The supernatant from these cells was added to Sanfilippo B patient fibroblasts in the presence or absence of mannose 6-phosphate. The study demonstrated uptake of NAGLU from the supernatant of these lentiviral-modified cells into Sanfilippo B fibroblasts that was nearly completely abolished with the addition of mannose 6-phosphate to the media, suggesting that NAGLU produced by gene therapy may not suffer the same post-translational processing issues as recombinant NAGLU enzyme produced by cells *in vitro*, as was used in this study.

In this study, as well as in our previous studies, enzyme replacement therapy administered into the cerebral ventricle improved heparan sulfate and beta-hexosaminidase activity levels in the liver and heart. The importance of treating disease outside of the brain in Sanfilippo B syndrome is not known. Cardiac involvement is described in patients, including valvular heart disease, myocardial thickening, and arrhythmia (*52, 53*). The disease course is progressive, and while currently cardiac involvement is not a driver of mortality or even major symptoms due to Sanfilippo B syndrome, it seems probable that if neurological disease is successfully treated any morbidity from heart disease would become more evident.

Cerebrospinal fluid bulk turnover occurs multiple times per day (approximately three times per day in humans), and so it is expected that recombinant enzymes administered into the cerebrospinal fluid would reach the blood circulatory system, albeit at a lower concentration, from which they would be taken up by organs and tissues outside the central nervous system. It is not known whether this approach of delivering enzymes to the cerebrospinal fluid in order to treat both the central nervous system and the rest of the body would be sufficient, holistic therapy for lysosomal diseases that affect body as well as brain.

Canine proteomic analysis revealed altered expression of lysosomal proteins, such as elevation of alpha-*N*-acetylglucosamine-6-sulfatase (GNS), cathepsin D (CTSD), and beta-hexosaminidase (HEXB). These results are not surprising, as lysosomal enzymes and other proteins may show elevated activity levels in tissues of Sanfilippo B mice compared to controls. Marked deficiency in the protein dipeptidyl peptidase 7 (DPP7), has not to our knowledge been previously associated with Sanfilippo B syndrome. Dipeptidyl peptidase 7 is a serine protease that localizes to non-lysosomal vesicles and shows substrate specificity similar to that of the cell-surface protein dipeptidyl peptidase 4 (*54*). Interestingly, HS and heparin oligosaccharides may inhibit the activity of dipeptidyl peptidase 4 (*55*). Low expression of collagen type I alpha1 (COL1A1), collagen type I alpha2 (COL1A2), and fibulin 5 (FBLN5), proteins implicated in connective tissue disorders, was found in Sanfilippo B dogs compared to controls, with more normal expression in treated animals. Deficiency in collagen and fibulin is not known to be associated with Sanfilippo B syndrome, but COL1A1 and COL1A2 have been described in the glycosaminoglycan interactome (*56*).

Our study had a number of important limitations. First, the rhNAGLU-IGF2 that was available for our studies was not materially the same as the candidate that is currently in clinical trials. We do not know the differences in composition between the clinical candidate material and the research grade enzyme that we used here. Second, our studies were not designed to evaluate the biodistribution of either rhNAGLU or rhNAGLU-IGF2 in the central nervous system. This is because our chosen study termination occurred four weeks after the final dose, following behavioral studies. The half-life of rhNAGLU-IGF2 in the brain was previously estimated at approximately seven days (*14*), although mice treated at postnatal day 1 or 2 showed elevated NAGLU activity at four weeks following a single ICV injection of rhNAGLU-IGF2 (*15*). However, the dose of enzyme (100 µg) was not increased as the mice grew (*15*), so that in this study little to no enzyme could be detected in the brain of these mice at four weeks after the final infusion. This limitation also meant that we were unable to determine the cellular distribution of the enzymes within tissues. Third, some behavioral assessments such as radial arm maze and social interaction test did not show intergroup differences between carriers and affected mice during this experiment, so that we could not use these tests to determine whether there was a therapeutic effect in treated cohorts. All the mice were cannulated, which may have affected their behavior and clearly limited the types of experiments we could conduct; for example, we were not able to perform Morris water maze testing in cannulated mice. Both control and treatment group experimental mice also underwent repeated sedation in order to dose vehicle or enzyme, and it was important to keep treatment as consistent as possible among groups. However, the repeated sedation events may have affected behavioral experiments and impacted overall survival, even in the otherwise healthy carrier mice.

## Conclusions

Long-term treatment with monthly intracerebroventricular rhNAGLU-IGF2 (research grade) showed a beneficial effect in Sanfilippo B mice. Significant intergroup differences were noted in survival, motor activity, stretch attend postures (fear behavior), and histological markers of microglial activation, and lysosomal substrate accumulation (HS, beta-hexosaminidase, LAMP1) between enzyme-treated and vehicle-treated Sanfilippo B mice. Whole tissue brain and heart HS levels were lower in Sanfilippo B mice treated with rhNAGLU compared to rhNAGLU-IGF2. We did not detect other intergroup differences between rhNAGLU and rhNAGLU-IGF2-treated mice in the outcome measures that we assessed in these experiments. Notably, our studies were not designed to assess NAGLU distribution, and only one experiment included treatment groups with both enzymes. Sanfilippo B dogs treated with rhNAGLU-IGF2 showed less HS in brain and CSF compared to untreated affected controls, with beta-hexosaminidase activity normalized to carrier levels in most cerebral areas including gray and white matter regions. Proteomics studies in cerebral cortex showed dysregulated pathways in Sanfilippo B dogs compared to carrier controls, with restoration of a more normal pattern in Sanfilippo B dogs treated with rhNAGLU-IGF2. These findings suggest that rhNAGLU-IGF2 administered into cerebrospinal fluid shows a beneficial effect on the Sanfilippo B phenotype in multiple species.

## Acknowledgements

P.I.D., J.T.D., M.S.S., and J.D.C. designed experiments and prepared the manuscript. S.Q.L., S.-h.K., M.S.R., and J.T.D. performed mouse experiments. S.Q.L, A.S., H.R.N., K.C., C.V. and J.D.C. performed histology including immunofluorescence and quantitative measurements. E.M.S., B.N.V., N.M.E., J.J., and J.D.S. performed canine experiments. F.W., S.L, S.Q.L., and T.F.C. performed enzyme assays and/or proteomics on canine tissues. We gratefully acknowledge the assistance and support of Soila Sukipolvi, Irina Zhuravka, George Lopez, Ling Wang, Valentina Sanguez, and Catalina Guerra. BioMarin Pharmaceutical and Allievex provided vehicle, rhNAGLU, and rhNAGLU-IGF2. BioMarin provided HS measurement of mouse brain and heart tissues. We gratefully acknowledge Roger Lawrence and Brett Crawford for their specific assistance with these assays.

