## Supplemental Figures and Tables S1-S3 for "Recombinant NAGLU-IGF2 prevents physical and neurological disease and improves survival in Sanfilippo B syndrome"

SUPPLEMENTAL MATERIAL

**Supplemental Table S1.** NAGLU Activity in Mice (28 days post-dose)

| | Brain 1+2 | Brain 3 | Brain 4 | Brain 5+6 | Spinal Cord | Liver | Heart |
|---|---|---|---|---|---|---|---|
| Mutant + Vehicle | 0.00 ± 0.00 | 0.00 ± 0.00 | 0.00 ± 0.00 | 0.00 ± 0.00 | 0.00 ± 0.00 | 0.00 ± 0.00 | 0.00 ± 0.00 |
| Carrier + Vehicle | 0.11 ± 0.01 | 0.08 ± 0.01 | 0.10 ± 0.01 | 0.10 ± 0.01 | 0.06 ± 0.02 | 0.55 ± 0.10 | 0.27 ± 0.05 |
| Mutant + NAGLU | 0.02 ± 0.01 | 0.03 ± 0.01 | 0.04 ± 0.02 | 0.02 ± 0.01 | 0.00 ± 0.00 | 0.00 ± 0.01 | 0.00 ± 0.00 |
| Mutant + NAG-IGF2 | 0.01 ± 0.01 | 0.03 ± 0.04 | 0.01 ± 0.01* | 0.01 ± 0.03 | 0.00 ± 0.00 | 0.00 ± 0.01 | 0.00 ± 0.00 |

Values are means ± standard deviations and are expressed as units of activity per milligram protein in homogenized tissue. Brains were divided coronally and numbered rostral to caudal 1-6, with the 1st and 2nd combined and the 5th and 6th combined for the assay. *outlier >2 S.D. excluded from analysis.

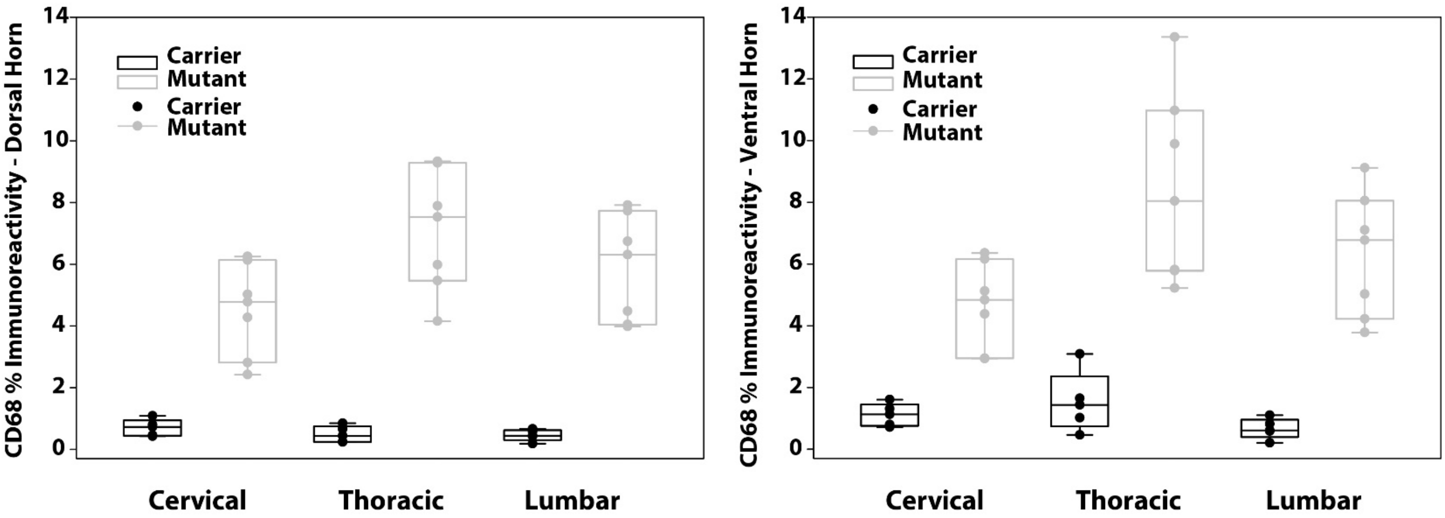


**Supplemental Figure S1**. Box plots of immunostaining for CD68 in cervical, thoracic, and lumbar spinal cord of Sanfilippo B (Mutant) and carrier mice. Boxes depict median, first and third quartiles, and range of immunoreactivity. Dots represent individual mice. $p \leq 0.001$ carrier v mutant in all regions shown.

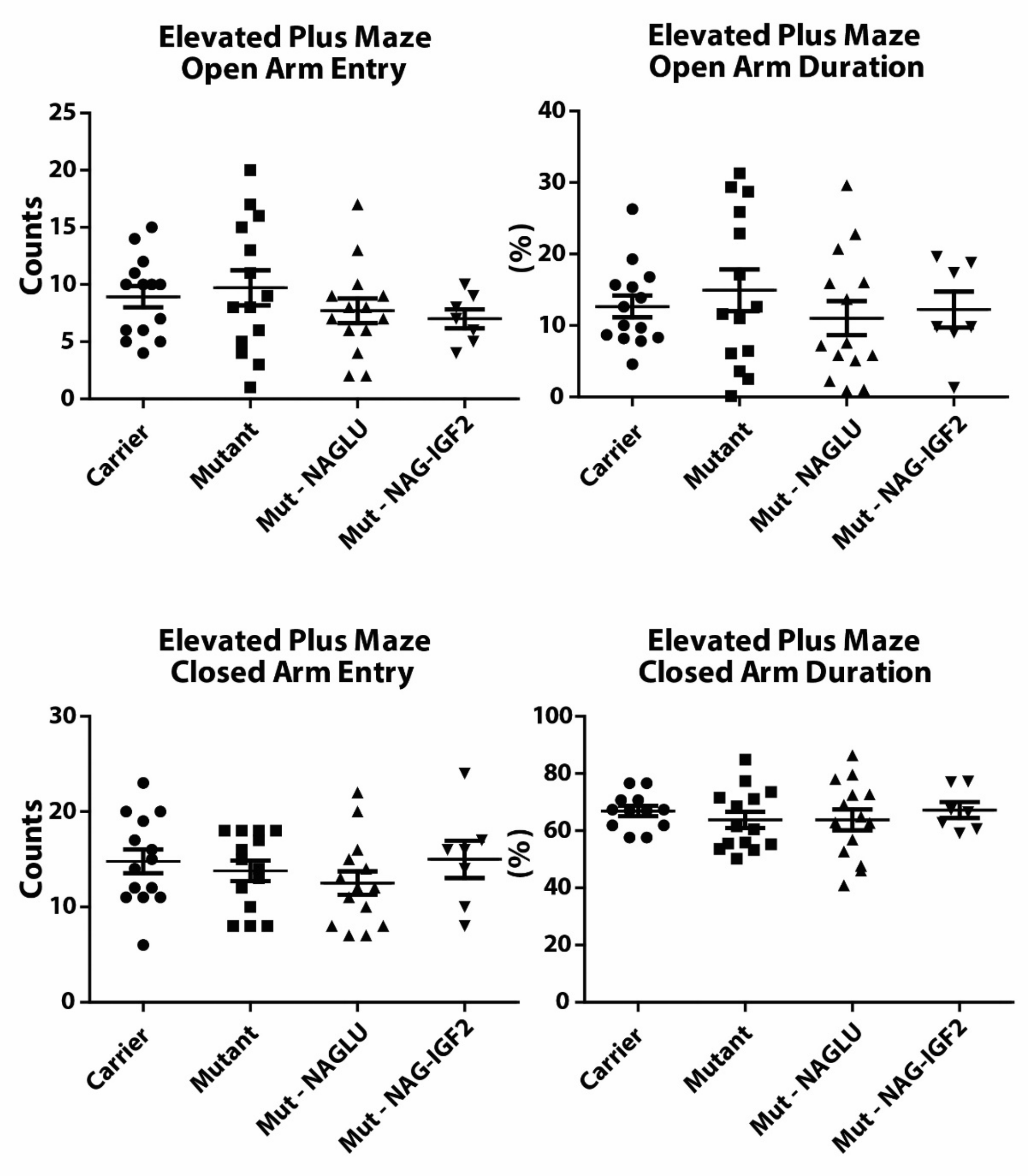


**Supplemental Figure S2.** Elevated plus maze. Symbols represent individual mice. Bars represent means and s.e.m. NAGLU: rhNAGLU. NAG-IGF2: rhNAGLU-IGF2. Mutant and Carrier control mice were treated with vehicle as described in the methods.

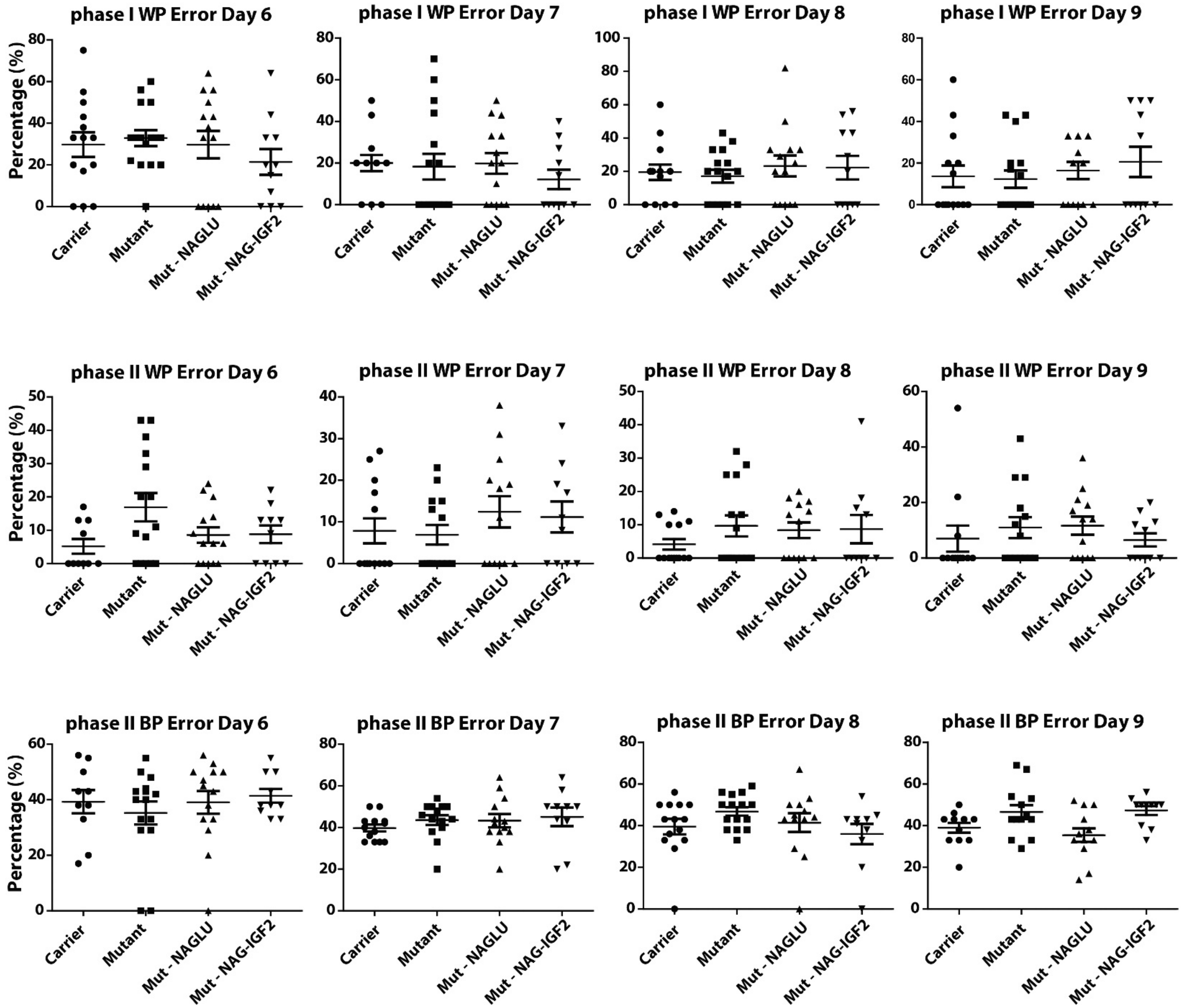


**Supplemental Figure S3.** Radial arm maze. WP: within-phase. BP: between-phase (long-term memory). Symbols represent individual mice. Bars represent means and s.e.m. NAGLU: rhNAGLU. NAG-IGF2: rhNAGLU-IGF2. Mutant and Carrier control mice were treated with vehicle as described in the methods.

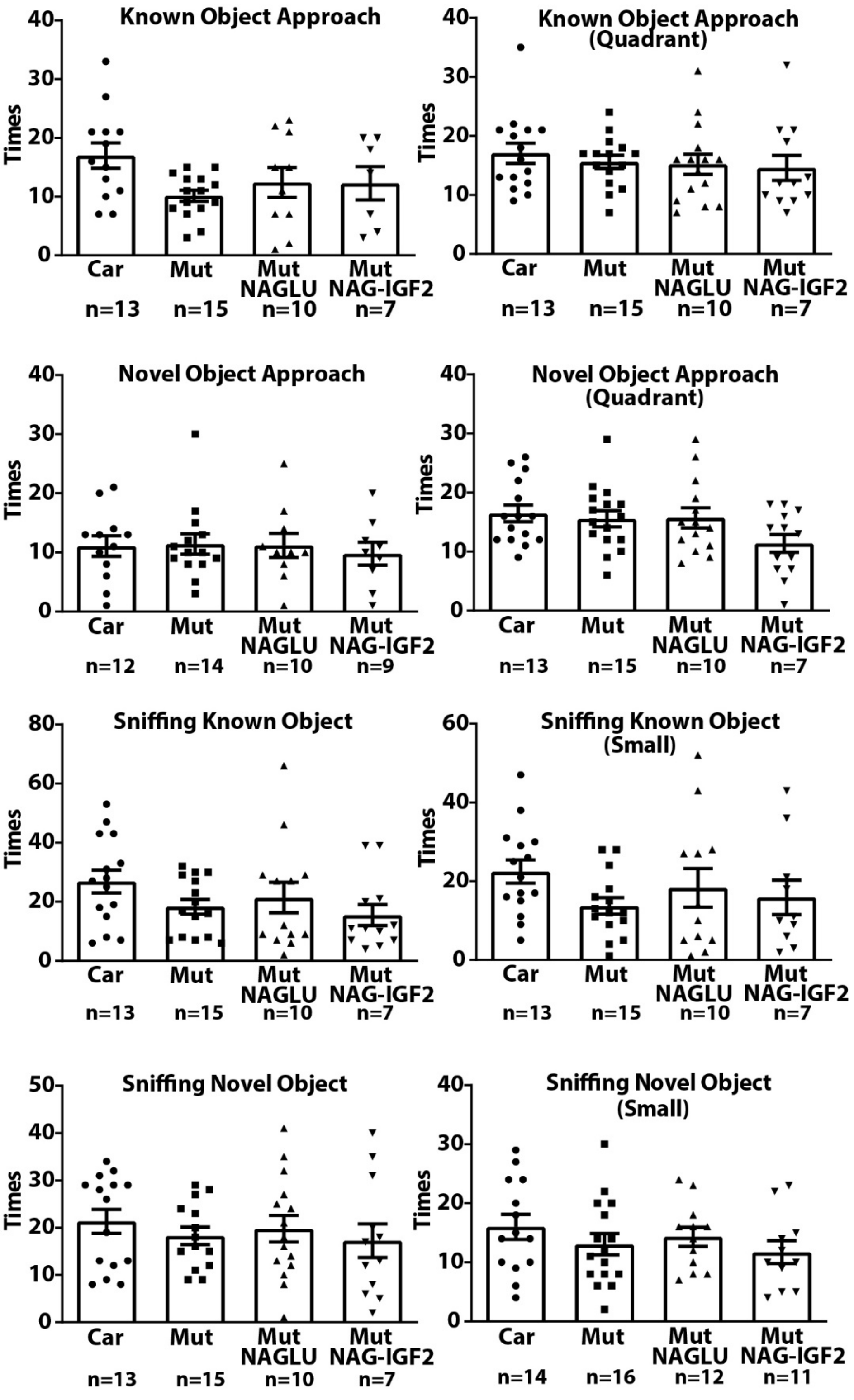


**Supplemental Figure S4.** Novel object recognition. Symbols represent individual mice. Bars represent means with s.e.m. NAGLU: rhNAGLU. NAG-IGF2: rhNAGLU-IGF2. Mutant and Carrier control mice were treated with vehicle as described in the methods.

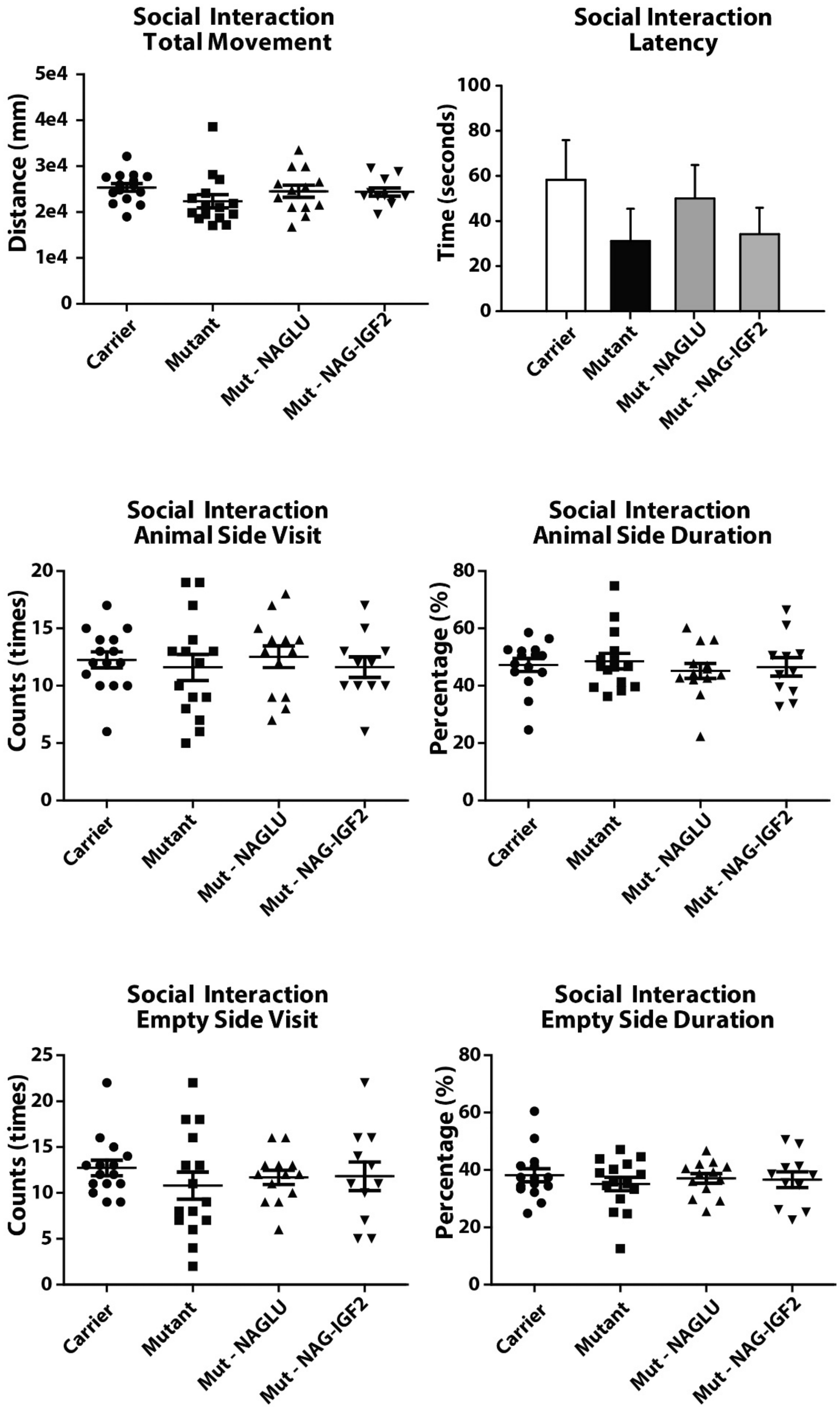


**Supplemental Figure S5.** Social interaction test. Symbols represent individual mice. Bars represent means and s.e.m. NAGLU: rhNAGLU. NAG-IGF2: rhNAGLU-IGF2. Mutant and Carrier control mice were treated with vehicle as described in the methods.

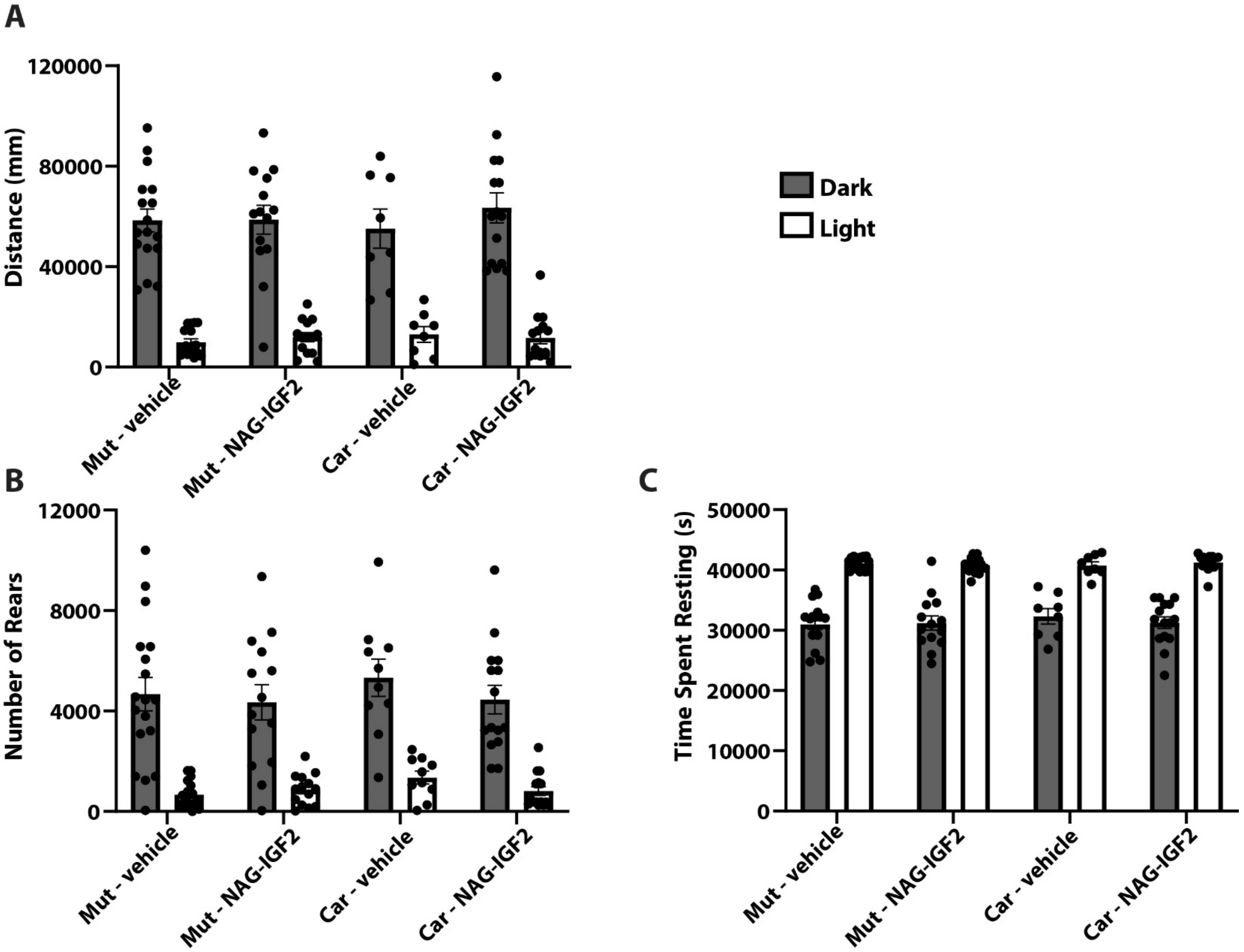


**Supplemental Figure S6.** Activity in dark and light phases of open field by genotype and treatment group at 22 weeks of age. **(A)** Overall distance traveled. **(B)** Number of rears. **(C)** Time spent resting. Mutant + vehicle, n=18. Mutant + NAG-IGF2, n=15. Carrier + vehicle, n=10. Carrier + NAG-IGF2, n=15. Means, standard error, and individual mice shown.

**Supplemental Table S2.** Survival study events (mice)

| Survival Study Events | Mutant + Vehicle | Mutant + NAG-IGF2 | Carrier + Vehicle | Carrier + NAG-IGF2 |
|---|---|---|---|---|
| Euthanized | 2F, 1M | 2F, 4M | 7F | 4F, 3M |
| Found dead | 3F, 1M | 3F, 4M | 2F, 1M | 1F, 1M |
| Died after Benadryl | 3F, 5M | N/A | N/A | 1F |
| Died after infusion | 1M | 2F | N/A | N/A |
| Taken down at 95 weeks | N/A | 1F | 4F, 2M | 2F, 4M |
| Cannula drop/ Recannulated | N/A | 3F, 1M | 5F, 1M | 2F, 3M |

M: male, F: female. N/A: not applicable

Euthanized: Euthanize to humane endpoints - weight loss, decrease appetite, weakness, moribund state
Found dead: Found dead in cage
Died after Benadryl: Found directly after intraperitoneal injection before ICV infusion
Died after infusion: Found directly after infusion
Taken down at 95 weeks
Cannula drop/Recannulated: n=10 recannulated once, n=5 recannulated multiple times (*N.B.,* these events are not fatalities)

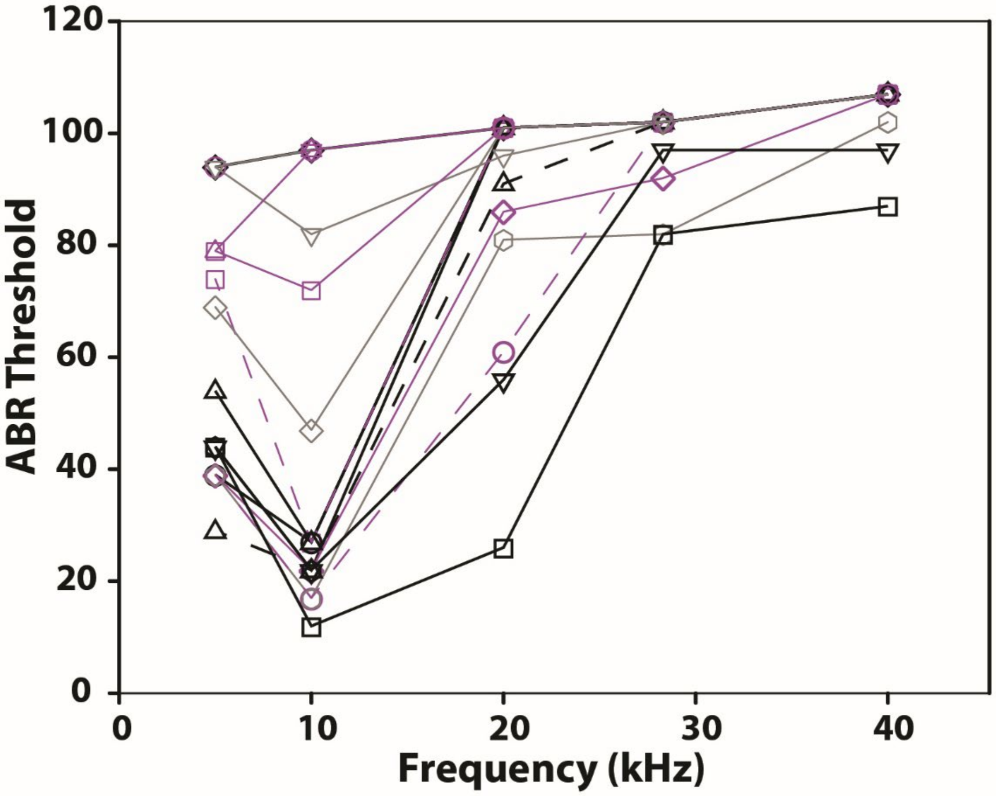


**Supplemental Figure S7.** Auditory brainstem responses (ABR) in mice. Individual mice are represented by symbols; connecting lines are added for clarity. Gray: Carrier mice treated with rhNAGLU-IGF2. Black: Carrier mice treated with vehicle. Purple: Sanfilippo B mice treated with rhNAGLU-IGF2.

**Supplemental Table S3.** NAGLU Activity in canine brain (four weeks post-dose)

| | Vermis | L-CSO | L-CN | L-CWM | R-CSO | R-CN | R-CGM | R-CWM |
|---|---|---|---|---|---|---|---|---|
| Mutant, Untreated | 0.00 ± 0.00 | 0.00 ± 0.00 | 0.00 ± 0.00 | 0.00 ± 0.00 | 0.00 ± 0.00 | 0.00 ± 0.00 | 0.00 ± 0.00 | 0.00 ± 0.00 |
| Carrier, Untreated | 0.467 ± 0.082 | 0.654 ± 0.096 | 0.512 ± 0.257 | 1.18 ± 0.236 | 0.516 ± 0.158 | 0.397 ± 0.250 | 0.397 ± 0.103 | 0.901 ± 0.089 |
| Mutant + NAG-IGF2 | 0.110 ± 0.129 | 0.00 ± 0.00 | 0.00 ± 0.00 | 0.00 ± 0.00 | 0.00 ± 0.00 | 0.00 ± 0.00 | 0.554 ± 0.513 | 0.00 ± 0.00 |

Values are means ± standard deviations and are expressed as units of activity per milligram protein in homogenized tissue. L: left, R: right, CSO: centrum semiovale, CN: caudate nucleus, CGM: cerebral gray matter, CWM: cerebral white matter.
